# Preclinical testing of RRx-001 in mouse models of experimental endometriosis reveals promising therapeutic impacts

**DOI:** 10.1101/2024.08.12.607591

**Authors:** Iona McIntyre, Vadim Vasilyev, Chiara Lia Perrone, Priya Dhami, Kavita Panir, Matthew Rosser, Erin Greaves

## Abstract

Endometriosis is a chronic inflammatory condition characterised by the presence of ectopic endometrial-like tissue (lesions), reduced fertility and chronic pain. Impacting both the health and psycho-social functioning of millions of women worldwide, there is an urgent need for innovative non-hormonal, non-invasive treatments for the disorder. Both peritoneal and lesion-resident macrophages have been strongly implicated in the pathogenesis of endometriosis; key roles include promotion of lesion growth, neuroangiogenesis and nerve sensitization. With such a central role in the disease, macrophages represent a novel therapeutic target. In the current preclinical study, we sought to repurpose the macrophage targeting anti-cancer drug RRx-001 for the treatment of endometriosis. We utilised mouse models of induced endometriosis to demonstrate that RRx-001 induces regression of endometriosis lesions and attenuates pain-like behaviours, without negatively impacting fertility. Targeted depletion of peritoneal cavity macrophages significantly impairs lesion reduction, demonstrating their critical role in mediating the anti-endometriosis impact of RRx-001. Using single nuclei multiomics, we identified a modification of macrophage subpopulations in the peritoneal cavity, specifically reduced acquisition of a pro-disease phenotype and an accumulation of a pro-resolving phenotype. These observations signify the potential of RRx-001 as a novel therapeutic for endometriosis management.

## Introduction

Endometriosis is a common, chronic, neuro-inflammatory condition that affects 1 in 10 women (and those assigned female sex at birth) during their reproductive years, although adolescent and postmenopausal cases are well documented(*1*). Endometriosis is defined by ectopic endometrial-like tissue which forms non-neoplastic ‘lesions’, commonly within the peritoneal cavity and rarely (<1%), in extra-pelvic locations (*2, 3*). Endometriosis has a complex and diverse symptomology which includes severe dysmenorrhea, chronic pelvic pain, dyspareunia, fatigue, and infertility. These symptoms have a cumulative impact on health, well-being, and psychosocial functioning. Beyond the personal impact on those that suffer from the condition and that of their partners and family, endometriosis also represents a substantial socioeconomic burden, due not only to the cost of treatment, but also due to significant reductions in productivity(*4*).

Despite the high prevalence of endometriosis, there remains limited etiological understanding of the disease(*5*). Sampsons’ theory of retrograde menstruation is the most commonly accepted hypothesis for the origin of endometrial-like tissue within the peritoneal cavity, referring to the reflux of sloughed menstrual debris through the fallopian tubes during menstruation(*6*). Although retrograde menstruation may be common, and reportedly occurring in up to 90% of those with patent fallopian tubes, only a proportion develop endometriosis, indicative of a multifactorial pathogenesis, with immune dysregulation prominent amongst many suspected etiological factors(*7–9*). Following an immunological inability to eliminate refluxed ectopic endometrial fragments, adherence and cell proliferation under hypoxic stress occurs, thus initiating the development of endometriotic lesions(*10, 11*). The current medical management of endometriosis is non-curative and includes analgesic management, hormonal suppression, and surgical debulking of lesions(*12*). This current strategy is insufficient due to common post-surgical recurrence of lesions, unwanted side-effects from hormonal medications (anovulation and those associated with hypoestrogenism), and a high prevalence of endometriosis-associated pain that is refractory to medical treatment(*13–15*). Thus, there is an urgent, unmet need for novel non-hormonal therapeutics for endometriosis.

Macrophages have been strongly implicated in the promotion of endometriosis, emerging as a potential target for a non-hormonal endometriosis therapy(*16*). These dynamic mononuclear phagocytes are present in all tissues, playing integral roles in innate immunity, tissue homeostasis, and the induction and resolution of sterile inflammation. Macrophages demonstrate considerable transcriptional and phenotypical plasticity, existing on a broad spectrum between two highly polarised states: pro-inflammatory and pro-repair, with a varied and dynamic functionality(*17*). Macrophage function is dependent on both the perception of extracellular signals within the tissue niche and cellular ontogeny: seeded pre-birth from the embryonic yolk sac/fetal liver, or monocyte-derived and recruited from the circulation(*18, 19*).

In endometriosis, macrophages (and their precursors, monocytes) are an important component of the intra- and extra-lesion microenvironment. They are continuously recruited and play pathophysiological roles in lesion establishment, angiogenesis, and nociception(*20, 21*). In both patients and animal models, pro-endometriosis macrophages have been demonstrated to trend predominantly towards a pro-repair phenotype(*22, 23*). However, broad heterogeneity in peritoneal and lesion-resident populations has been revealed in detailed analyses using techniques such as single cell RNA-sequencing (scRNA-seq) and mass cytometry by time-of-flight (mass CyTOF), demonstrating subpopulations across the spectrum, playing subtype-specific roles in the progression or regression of endometriosis(*24–27*). An inability of macrophages to sufficiently recognise and clear refluxed ectopic endometrial-like tissue, alongside deficient phagocytic capabilities, have been identified in several studies(*28*). In the peritoneal cavity of mice, the predominant macrophage populations are small peritoneal macrophages (SpM) and large peritoneal macrophages (LpM), with monocyte-derived LpM demonstrated to be protective against endometriosis development, possibly due to enhanced lipid metabolism and cholesterol efflux, highlighting an intrinsic link between macrophage ontogeny and functionality(*26, 29*). Conversely, endometrial-derived lesion-resident macrophages enhance lesion growth(*26, 29*) and monocytes that are recruited directly to lesions from the vasculature differentiate *in situ* into macrophages that exhibit a pro-disease phenotype(*26*). Macrophages and monocyte subpopulations are also altered in the endometrium and peripheral blood of women with endometriosis, suggesting systemic myeloid dysfunction(*30*).

Parallels in phenotype and function can be drawn between macrophages in endometriosis and solid cancers. As with lesions, monocytes are continuously recruited into tumour tissues where they differentiate into macrophages. Niche specific factors, including hypoxic stress, lead to phenotype specification and pro-disease functions and an attenuation of phagocytic capabilities(*31*). Tumour associated macrophages (TAM) are broadly described as having a pro-repair phenotype, however, similar to endometriosis macrophages, deep immunophenotyping has identified diverse TAM subpopulations across the phenotypic continuum, including macrophages in highly phagocytic, cytotoxic, and potentially anti-tumorigenic states(*32–34*). Paradoxically, when appropriately polarised, tumour-resident macrophages can mediate cancer cell phagocytosis, and enhance tumour clearance(*35*). Multiple anti-cancer immunotherapies that reprogram TAMs have emerged in recent years, aiming to alter pro-tumorigenic phenotypes and enhance tumour-clearing activity(*36*). Due to the inherent similarities between the immunopathophysiology, and recent single cell analysis revealing the presence of pro-disease TAM-like macrophages within endometriotic lesions(*26*), we have sought to repurpose the therapeutic strategy of macrophage repolarisation for endometriosis.

RRx-001 is a first-in-class small molecule inhibitor currently in several phase I-III clinical trials for a variety of solid cancers, with properties linked to multiple mechanisms of activity including macrophage modulation, phagocytic enhancement, vascular normalisation, inflammasome inhibition and epigenetic modifications, amongst others(*37*). In the current study, we have evaluated the impact of RRx-001 in preclinical mouse models of experimental endometriosis to determine the effect on endometriotic lesion growth and associated pain-like behaviours, and to identify any impacts on oestrous cyclicity and reproductive outcomes. As RRx-001 is known to modulate macrophage phenotype, we investigated the impact of targeted peritoneal cavity macrophage depletion and utilised 10x multiomic sequencing to investigate the transcriptomic and epigenetic landscape of peritoneal macrophages isolated from mice with induced endometriosis, dosed with or without RRx-001. Collectively, we describe vital preclinical data that indicates RRx-001 could be a promising new therapeutic for endometriosis

## Results

### RRx-001 reduced endometriotic lesion growth and attenuated pain-like behaviours in a ‘menses’ mouse model of experimental endometriosis

Initially, we assessed the therapeutic potential of RRx-001 for endometriosis treatment, utilising a pre-treatment approach in a syngeneic bioluminescent ‘menses’ mouse model of induced endometriosis. As macrophages are essential in the early establishment of lesions, a pre-treatment approach was used to evaluate whether RRx-001 altered lesion initiation or subsequent progression. In brief, ovariectomised recipient mice received an intraperitoneal injection of ‘menses’-like endometrium and oestradiol supplementation(*38*). Mice were randomly assigned to a treatment group and received either intravenous injection of RRx-001 or vehicle control, 6 hours prior to donor tissue transfer, and then twice weekly for 2 weeks (Fig.1A). *In vivo* bioluminescent imaging at days 3, 7, 10 and 14 was performed for continuous monitoring of lesions. In both the vehicle control and RRx-001 treated groups, lesions were detectable from day 3 onwards, demonstrating that RRx-001 does not prevent initial lesion development (Figs. 1B, 1C). Some spontaneous resolution and/or lesion progression was observed in the ‘menses’ model of endometriosis (as previously documented(*39*)), leading to high intragroup variability of bioluminescent signal(*38*). Despite the occurrence of some spontaneous resolution, there was no significant difference in bioluminescent signal across the vehicle group at different timepoints. Conversely, mice dosed with RRx-001 exhibited a reduction between days 3 and 7, and days 3 and 14 (p=0.0079 and p=0.020, respectively; Fig.1C). Lesions were recovered at day 14 and histologically verified (Fig.S1); mice dosed with RRx-001 exhibited decreased lesion number (p=0.013; Fig.1D) and a reduced cross-sectional lesion area (median = 1.48 mm^2^), compared to vehicle control (median = 4 mm^2^, p=0.04; Fig.1E). To determine if RRx-001 attenuated evoked pain-like behaviours, mechanical hyperalgesia was assessed using von Frey filaments on days 11, 12 and 13 post-lesion induction. Mice that received RRx-001 exhibited an increased abdominal retraction threshold (p=0.0004; Fig.1F), implying reduced abdominal sensitivity, possibly due to the resolution of lesions within the peritoneal cavity. No differences were observed in hind paw withdrawal measurements (Fig.1G).

**Figure 1.**
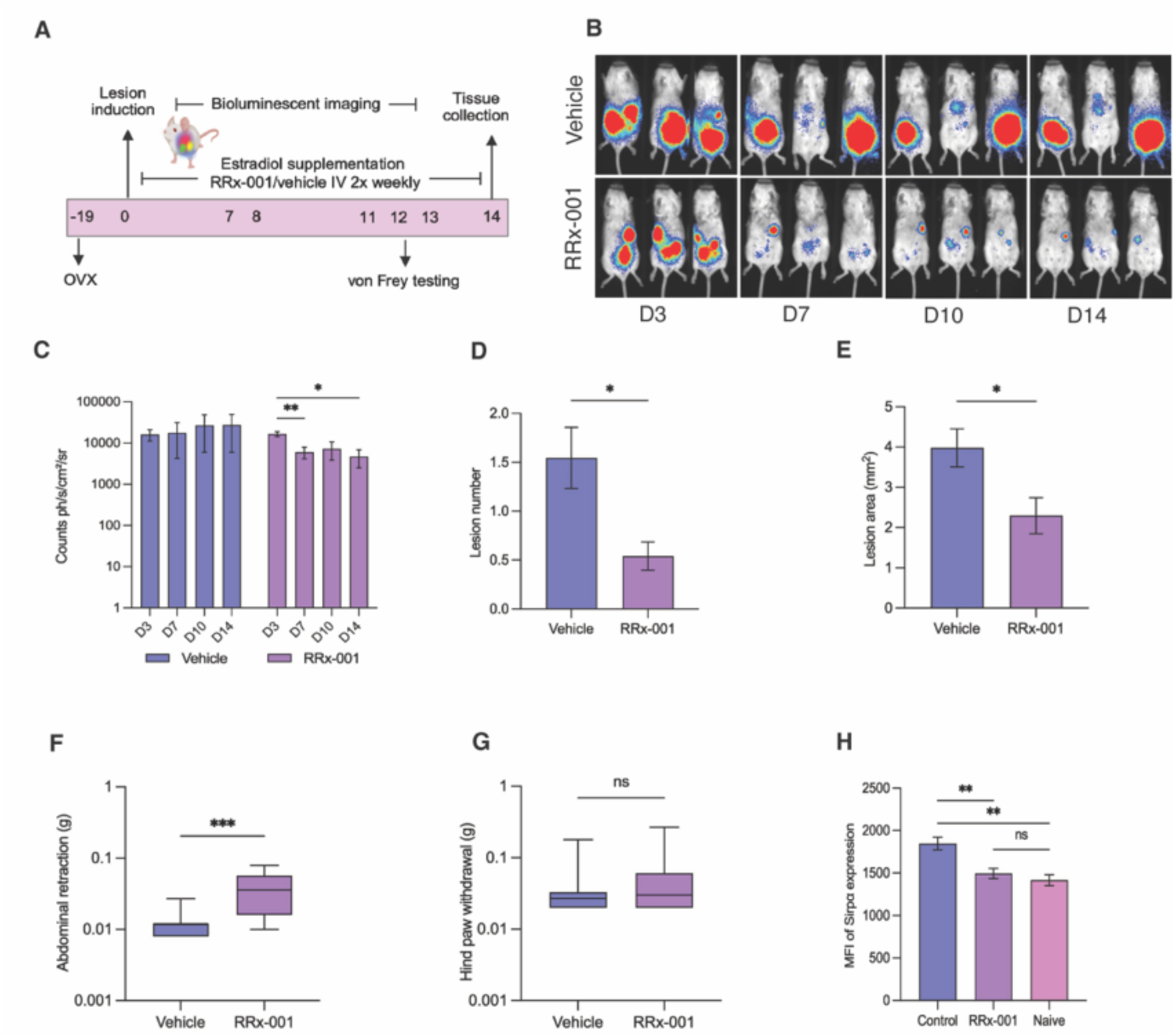
RRx-001 reduces lesion growth and mechanical allodynia in a ‘menses’ model of experimental endometriosis. (A) Schematic of experimental design. (B) Representative images from days 3, 7, 10 and 14 from RRx-001 and vehicle treated mice. (C) Quantification of lesion bioluminescent signal of RRx-001 and vehicle treated mice with endometriosis; statistical analysis performed using a two-way ANOVA with Bonferroni post hoc. (D) Number of lesions and (E) size of lesions recovered from RRx-001 and vehicle mice; statistical analyses performed using a Mann-Whitney test. (F) Mechanical hyperalgesia measured using von Frey filaments applied to the abdomen and (G) hind paw (F and G plotted as an average of 3 measurements taken over days 11-13 post-endometriosis induction). Statistical analyses performed using a Mann-Whitney test. (H) Median fluorescence intensity (MFI) of Sirpα on large peritoneal macrophages; statistical analysis performed using a one-way ANOVA. * p < 0.05, ** p < 0.01, **** p < 0.001.

We performed flow cytometry analysis on the peritoneal lavage of endometriosis mice treated with or without RRx-001 and found no alteration in the abundance of LpM or SpM populations (Fig.S2A and C-D, gating strategy in Fig.S2F). As RRx-001 has been reported to modulate macrophage phenotype, and was previously found to attenuate expression of Sirpα(*40*) (receptor Sirpα-Cd47 ligand interaction protects ‘self’ cells from phagocytosis), we investigated the expression of Sirpα on peritoneal macrophages; expression was increased on LpM isolated from mice with induced endometriosis when compared with naïve mice (no procedures) (p = 0.0046, Fig.S2B and E) and RRx-001 treatment attenuated Sirpα levels (p=0.0053; Fig.S1H).

### Depletion of macrophages by clodronate compromises the efficacy of RRx-001

As our initial findings indicated that treatment with RRx-001 led to lesion clearance and phenotypical modulation of peritoneal macrophages, but did not prevent lesion establishment, we next assessed the therapeutic impact of RRx-001 in a model of established endometriotic lesions with concomitant macrophage depletion, to confirm that the anti-endometriosis impact seen was driven by macrophages, as previously described in anticancer studies (*58*). Mice were randomly assigned to a treatment group (I) RRx-001 or (II) RRx-001+clodronate and then received an injection of ‘menses-like’ endometrium. Following a one-week period of lesion establishment, mice received 3 intravenous doses of RRx-001, alongside intraperitoneal liposomal clodronate for targeted macrophage depletion in the RRx-001+clodronate group (Fig.2A). Depletion of peritoneal macrophages in the RRx-001+clodronate group was confirmed using flow cytometry analysis, with the proportion of peritoneal macrophages significantly reduced compared to the RRx-001 group (p= 0.0286, Figs.2B, 2C). *In vivo* bioluminescent imaging was performed at day 7 (pre-treatment) and day 14 (post-treatment) (Fig.2D). No difference in lesion bioluminescence was detected between groups at day 7 (Fig.2E). At day 14, following treatment, a significant difference in lesion bioluminescence was observed between treatment groups (p=0.0173, Fig.2E), with mice treated with RRx-001 alone exhibiting reduced lesion bioluminescence, and the RRx-001+clodronate mice exhibiting increased lesion bioluminescence, suggesting a critical role for macrophages in mediating the therapeutic effect of RRx-001.

**Figure 2.**
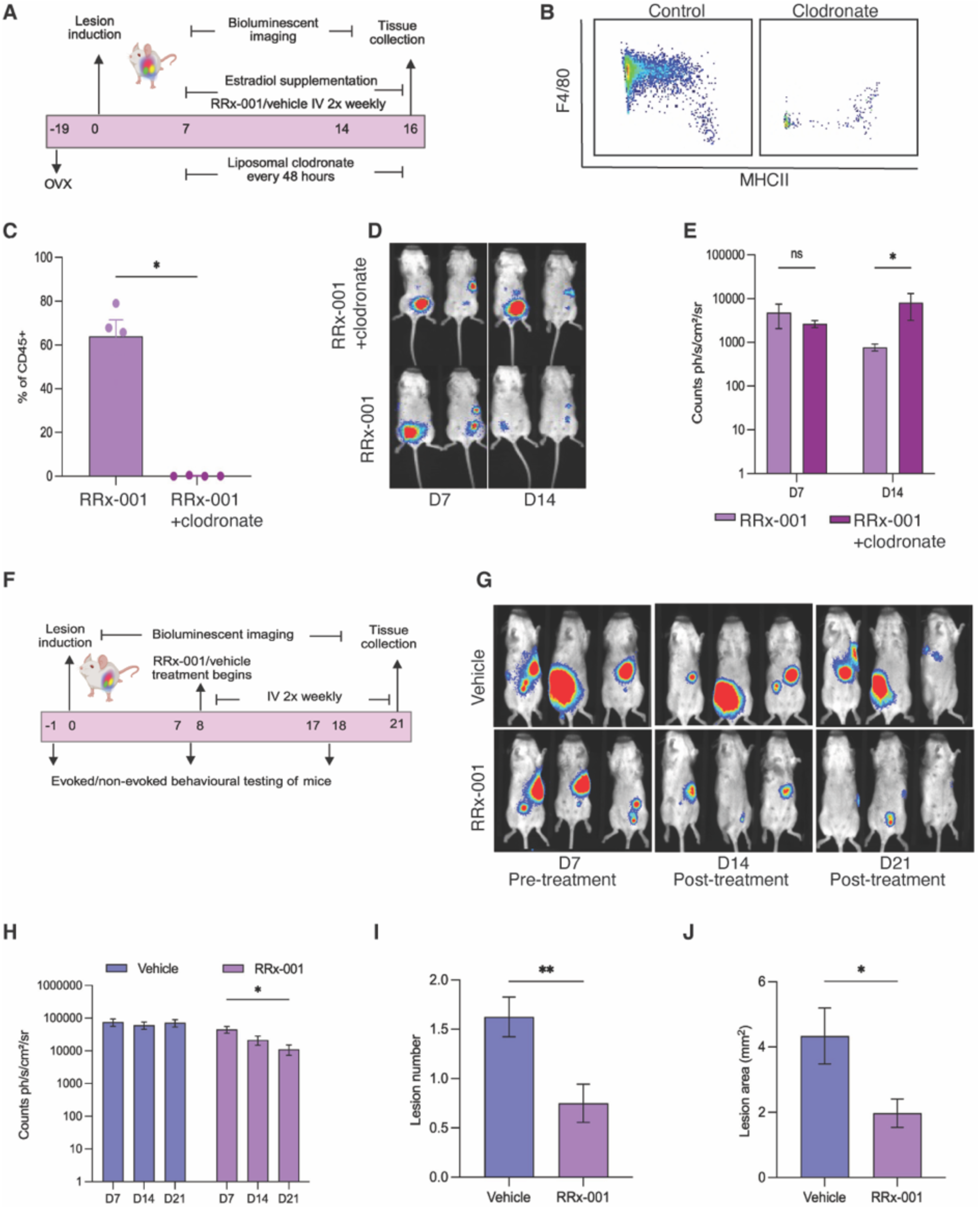
The macrophage-mediated therapeutic impact of RRx-001 on established endometriotic lesions in the menses and minimally invasive model of induced endometriosis. (A) Schematic of experimental design for combined RRx-001 treatment and clodronate depletion in the ‘menses’ model. (B) Representative flow cytometry plots of F4/80^+^ peritoneal macrophages from RRx-001 and RRx-001+clodronate treated mice, gated as in Fig.S2E. (C) Percentage of F4/80^+^ macrophages in the Lin^-^ CD45^+^ cells. (D) Representative images from days 7 and 14 from RRx-001 and RRx-001+clodronate treated mice. (E) Quantification of bioluminescent signal of RRx-001 and RRx-001+clodronate treated mice; statistical analysis performed using multiple Mann-Whitney tests with Bonferroni-Dunn correction for multiple comparisons. (F) Schematic of experimental design for post-treatment in the minimally invasive model. (G) Representative images from days 7, 14 and 21 from RRx-001 and vehicle treated mice. (H) Quantification of bioluminescent signal of RRx-001 and vehicle treated mice; statistical analysis performed using a two-way ANOVA with Bonferroni post hoc. (I) Number of lesions and (J) size of lesions recovered from RRx-001 and vehicle mice; statistical analyses performed using a Mann-Whitney test. * p < 0.05, ** p < 0.01.

### RRx-001 reduced endometriotic lesion growth and attenuated evoked *and* non-evoked pain-like behaviours in a minimally invasive mouse model of experimental endometriosis

To extend our pre-clinical testing of RRx-001, we utilised an additional, complementary mouse model of endometriosis. In this minimally invasive bioluminescent model, full thickness uterine fragments were injected intraperitoneally into intact recipient mice. Lesions established in this model exhibit less spontaneous regression and improved longevity(*38*), and since recipients are intact, post-incision pain sensitivity is avoided allowing improved pain behaviour testing (*41, 42*). Mice received a full thickness tissue transfer (day 0); one week following lesion induction, mice were dosed (day 8) with either RRx-001 or vehicle, according to randomly assigned treatment groups, and then twice weekly for 2 weeks (Fig.2F). Mice exposed to RRx-001 exhibited a reduction in lesion bioluminescence at day 21 compared to day 7 (pre-treatment) (p=0.25; Figs.2G, 2H). No reduction in bioluminescence was observed in the vehicle group. Lesions were recovered at day 21 and histologically verified. The RRx-001 group fewer lesions (p=0.0056; Fig.2I) and a reduced cross-sectional area (median = 1.6 mm^2^) compared to the vehicle treated mice (median = 3.1 mm^2^, p=0.020; Fig.2J).

To perform a comprehensive investigation into the impact of RRx-001 on endometriosis-associated pain, we used multiple evoked and non-evoked behavioural tests at i) baseline (pre-endometriosis induction), ii) after endometriosis induction but prior to treatment (days 7/8) and iii) after RRx-001/vehicle treatment (days 17/18; Fig.2A). To determine if RRx-001 treatment led to an attenuation of evoked pain-like behaviours, mechanical and thermal allodynia were assessed using von Frey filaments and the hot plate test, respectively. We did not detect any significant differences between groups prior to endometriosis induction (day −1/-2) or pre-treatment (day 7/8) across evoked behavioural tests (Fig.3A-C). Post treatment, RRx-001 treatment led to a significant decrease in mechanical allodynia, with increased withdrawal thresholds in both abdominal (p=0.002; Fig.3A) and hind paw measurements (p=0.013; Fig.3B). A significant increase in hot plate latency at day 17 in RRx-001 treated mice (p=0.0064; Fig.3C) was also detected. Next, we assessed the impact of RRx-001 on non-evoked pain-like behaviour using thermal plate preference and the ethologically relevant measures, burrowing and nesting. Burrowing and nesting are known to be reduced or eliminated with decreased well-being and can therefore be used as a surrogate marker of visceral pain evoked by endometriosis(*43*). Prior to endometriosis induction or pre-treatment at days 7/8, we did not detect any statistical differences between RRx-001 and vehicle groups in non-evoked behavioural tests (Fig.3D-F). Post treatment, thermal plate preference testing (TPPT) revealed a significant difference between groups at day 18 (p=0.005; Fig.3D) which may be indicative of a potential alleviation of hypothermal sensitivities following RRx-001 treatment. Of note, mice exhibited only exploratory behaviour on the cold plate, whereas the warm plate was also used for grooming by all mice, suggesting a thermal preference for warmth, as opposed to cold allodynia. No post-treatment differences were identified in burrowing behaviour between experimental groups (Fig.3E). RRx-001 treated mice did exhibit significantly increased nest complexity score (p<0.0001, Fig.3F, representative nest images shown in Fig.3G) indicating improved well-being due to an attenuation of endometriosis-associated pain.

**Figure 3.**
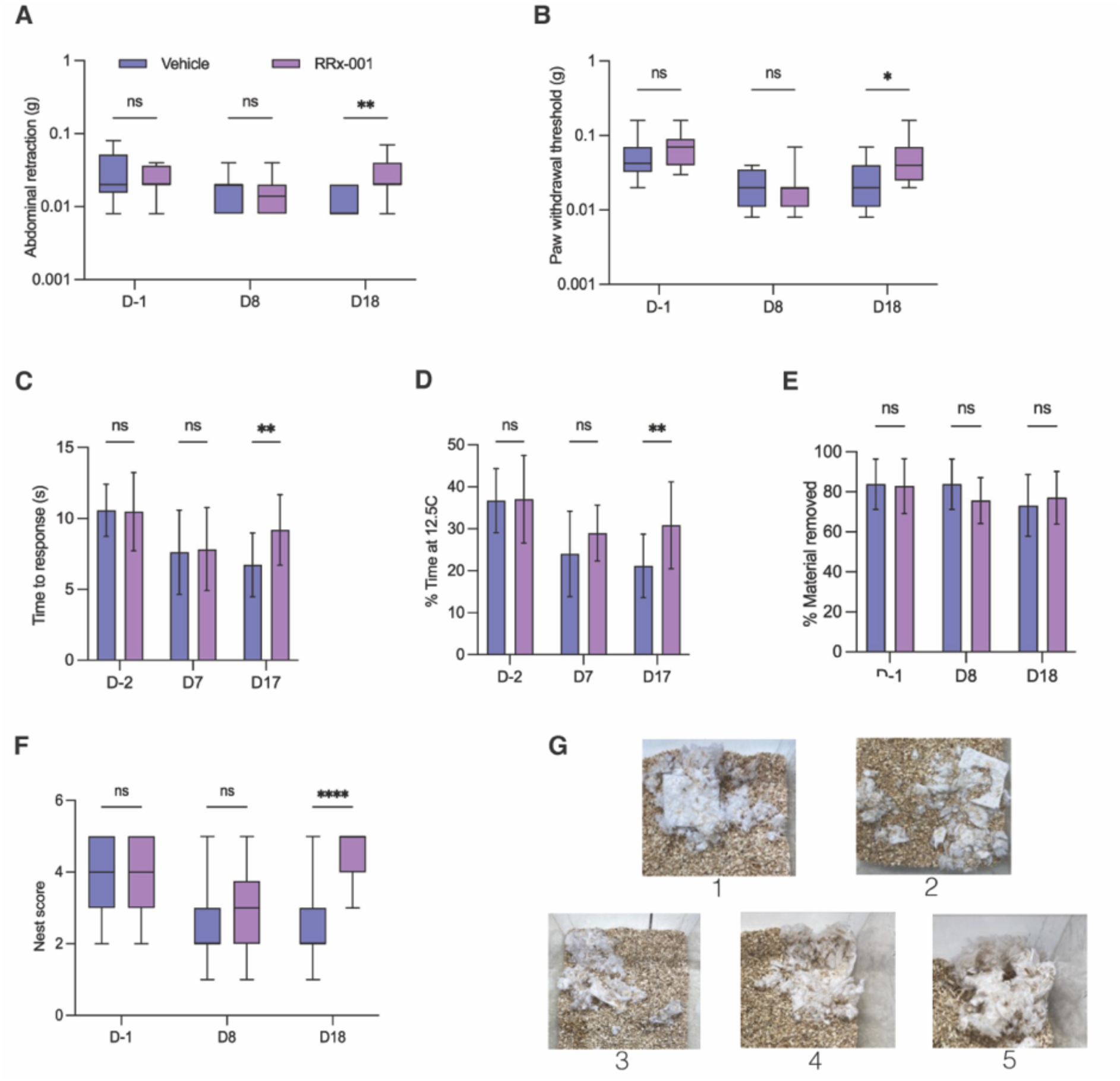
RRx-001 reduces evoked and non-evoked pain-associated behaviours in a minimally invasive model of induced endometriosis. (A) Mechanical allodynia measured using von Frey filaments applied to the abdomen and (B) hind paw; statistical analyses performed using multiple Mann-Whitney tests. (C) Hot plate latency assessed at 52°C; statistical analyses performed using multiple unpaired t-tests. (D) Percentage of material displaced in a burrowing assay; statistical analyses performed using multiple unpaired t-tests. (E) Thermal plate preference test analysis of % time spent on the 12.5 °C plate; statistical analyses performed using multiple unpaired t-tests. (F) Nest complexity scores; statistical analyses performed using multiple unpaired t-tests. (G) Example nests corresponding to scores 1-5. * p < 0.05, ** p < 0.01, **** p < 0.0001.

### RRx-001 impacts the transcriptomic and epigenomic landscape of peritoneal macrophages in mice with induced endometriosis

10x single nuclei multiomic profiling was used to generate paired RNA and ATAC libraries for sequencing to investigate the transcriptomic and epigenomic impacts of RRx-001 on peritoneal macrophages (Fig.4A). For this experiment, we used the ‘menses’ model of induced endometriosis, as the intraperitoneal injection of decidualised endometrial tissue most closely recapitulates the refluxed menstrual tissue thought to initiate endometriotic lesions. At day 7 post-lesion induction, fluorescent activated cell sorting (FACs) was utilised to isolate lineage-, Cd45+ leukocytes from the peritoneal lavage of RRx-001 and vehicle treated endometriosis mice, as well as sham controls (n=9, n=5, n=3, respectively, Fig.S3A), followed by multiomic processing using the 10X pipeline. We performed an integrative analysis of peritoneal macrophages combining both modalities into a ‘weighted-nearest-neighbour’ (WNN) framework, with any cells identified as non-macrophages removed (Fig.S3B-E)(*44*). UMAP projection revealed 7 macrophage clusters; inset shows the samples split by library ID (Fig.4B). We found consistent weighting of RNA and ATAC across clusters (although an increased RNA modality weight was seen in the proliferative cluster), uniformity of high-quality nucleosome signal (ATAC) and a consistently high log10GenesPerUMI (RNA) (Fig.S3C). All peritoneal macrophage clusters were *Cd45*+ and *Cd11b*+, and feature plots of *H2-Ab1* and *Adgre1* enable visualisation of SpM by low *Adgre1* (F4/80) and high *H2-Ab1* (MHCII) expression, prototypical LpM with high *Adgre1* and low *H2-Ab1*, and intermediate *Adgre1* in other LpM populations, as previously characterised in mice with induced endometriosis at the single cell level(*26, 45*) (Fig.4C).

**Figure 4.**
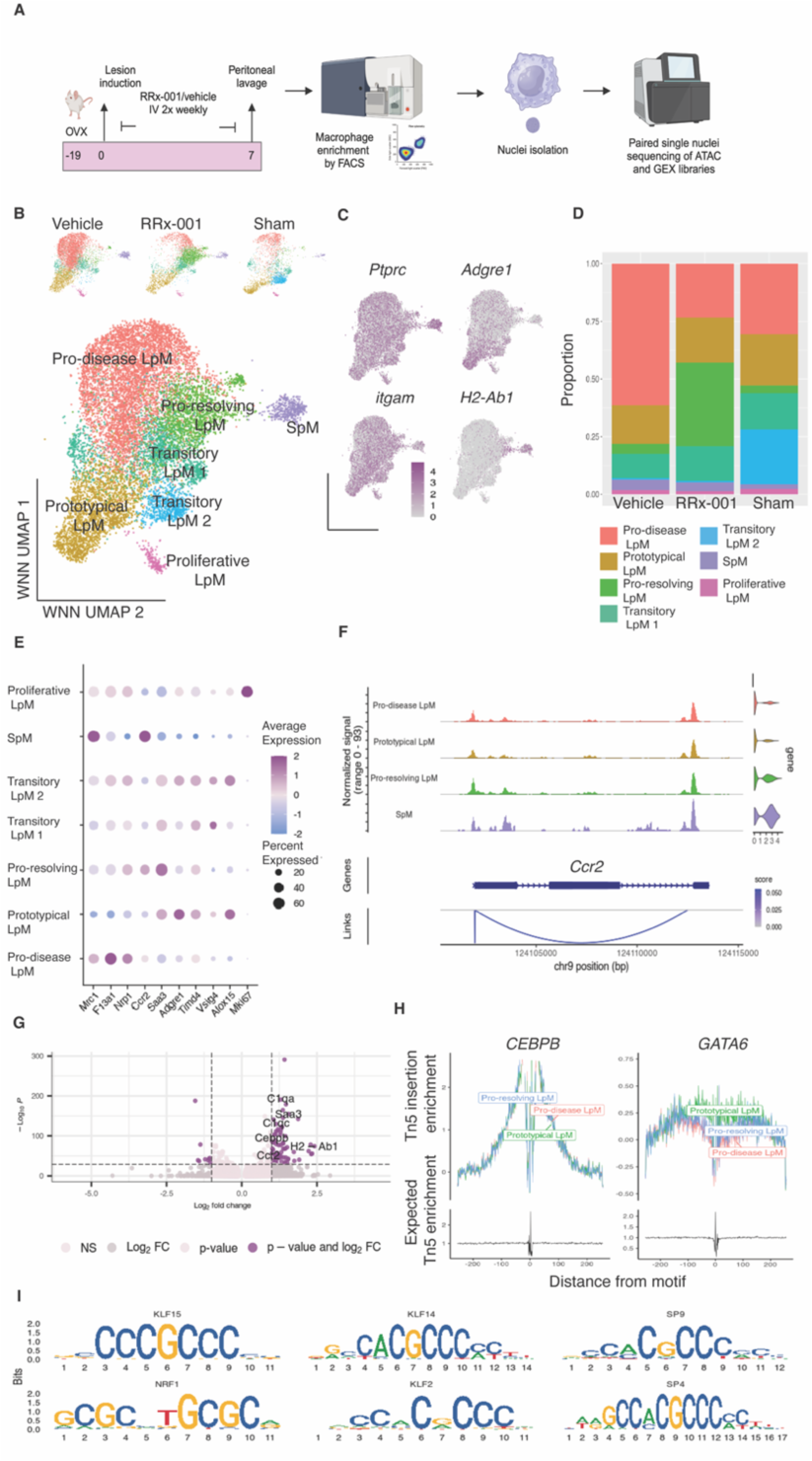
Multiomic analyses of RNA and ATAC modalities of peritoneal macrophages. (A) Schematic of experimental design. (B) UMAP visualisation based on a weighted combination of RNA and ATAC data from peritoneal macrophages isolated from vehicle treated endometriosis mice, RRx-001 treated endometriosis mice and sham mice. Inset is UMAP projection split by sample ID. (C) Feature plots of *Ptprc* (Cd45), *Itgam* (Cd11b), *Adgre1* (F4/80) and *H2-Ab1* (MHChcII) expression levels. (D) Samples split by cluster composition. (E) Dot plot of featured genes. (F) Chromatin accessibility, gene expression data and peak-gene links for Ccr2 expression in selected macrophage populations. (G) Volcano plot of up- and down-regulated DEGs of pro-resolving LpM compared to pro-disease LpM. (H) Transcription factor footprinting of Cebpb and Gata6 in pro-resolving LpM, pro-disease LpM and prototypical LpM populations. (I) DNA sequence motif enrichment analysis of ATAC data in peritoneal macrophages derived from RRx-001 treated endometriosis mice.

An analysis of sample composition by cluster revealed macrophage subpopulations were shared between experimental groups. Prototypical LpM (defined by *Timd4* and characteristic DEGs *Prg4*, *Ltbp1*, *Alox15*) exhibited comparable proportions in each sample, as did proliferative LpM and SpM. The predominant population in the vehicle treated endometriosis mice, termed pro-disease LpM, expressed the DEGs *F13a1, Gas6, Lyve1* and *Nrp1*, previously linked to endometriosis(*30, 45*), this population was markedly reduced in both RRx-001 treated and sham mice (Fig.4D,E). In RRx-001 treated mice, the predominant population was termed pro-resolving LpM due to the associated lesion resolution observed and the expression of genes such as *Ms4a4*, *Ccr2*, *Calhm6*, *Fcrla* and *Saa3* associated with inflammatory monocyte-derived infiltration, response to pathogens / antigens and lipid-associated phenotypes, previously linked to lesion resolution (Fig.4D,E)(*26, 29*). We included sham mice in this analysis, which exhibit a post-surgical loss and subsequent replenishment of prototypical LpM and therefore model a resolution of transient inflammation. As expected, transitory LpM were predominant in sham mice; both populations expressed intermediate levels of *Adgre1* and *Timd4* and were separated by enhanced *Vsig4* expression (in transitory LpM 1; Fig.4D,E), a gene known to represent longer time-in-residency, suggesting transitory LpM1 have been present in the peritoneal niche longer than transitory LpM2(*46*).

Abundant expression of *Ccr2* and low *Timd4* on SpM and pro-resolving LpM indicated recent recruitment/monocytic-derivation of these populations, an ontogeny previously shown to be protective against endometriosis development(*29*); paired ATAC data also demonstrated increased chromatin accessibility of the *Ccr2* gene in these populations and significant peak-gene links (Fig.4F). Top DEGs separating pro-resolving and pro-disease LpM were those linked with monocytic ontogeny and complement protein C1q as well as *Saa3* (Fig.4G). Enhanced mRNA levels of the transcription factor, *Cebpb* (associated with regulating macrophage lipid metabolism and phagocytosis(*47*)) were evident in the pro-resolving population (Fig.4G), suggesting enhanced Cebpb regulation of this phenotype. Conversely, transcription factor footprinting did not support this; with comparable enrichment observed in pro-resolving, pro-disease and prototypical populations. Expected footprinting differences, such as the characteristic Gata6 control of prototypical LpM, were detectable in the ATAC data, with most enrichment observed in prototypical LpM and least enrichment observed in pro-disease LpM (Fig.4H)(*48*). We also performed DNA sequence motif enrichment analysis of the ATAC data to identify overrepresented motifs in the differentially accessible peaks of RRx-001 treated mice (all macrophages), with hypergeometric probability testing of enrichment against a set of background peaks matched for GC content. We identified a strong enrichment of Kruppel-like factor motifs (Klf15, Klf14, Klf2), with Klf family members having broadly reported roles in the activation and polarisation of macrophages, recent research has identified highly specific and unexpected roles in macrophage differentiation for these broadly expressed transcription factors(*49*). The greatest overrepresentation was seen in the Klf14 (4.03-fold enriched, p < 0.0001) and Klf15 (2.91-fold enriched, p < 0.0001) motifs, which alongside Klf2, have been linked to modulation of inflammatory signalling pathways. Sp4 and 9 motifs were also enriched (3.06-fold and 2.96-fold enrichment respectively). Of particular interest was the enrichment of Nrf1 motifs (5.29-fold enrichment, p<0.0001); RRx-001 has previously been shown to induce Nrf2 which mediates an antioxidant response(*50*). Nrf1 works in concert with other transcription factors such as Nrf2 to modulate cellular responses to oxidative stress and maintain cellular homeostasis.

Next, slingshot was used to perform cell lineage and pseudotime inference analysis on reduced dimensional data using previously identified WNN clusters. SpM was specified as the initial cluster as the infiltration and transformation of monocytes into SpM with eventual differentiation into LpM is known to occur under inflammatory conditions, such as endometriosis. Lineage inference identified three branching trajectories from SpM (Fig.5A)(*23*); pseudo-temporal trajectories to the terminal populations of pro-disease LpM, prototypical LpM and proliferative LpM were identified. Interestingly, pro-resolving LpM appeared to be an intermediate in all trajectories, further supportive of a recent monocyte-derived ontogeny for this population (Fig.5B). Multinomial logistic regression was used to assess the pseudo-temporal sample imbalance in individual lineages, while adjusting for baseline sample probabilities. Macrophages from sham mice were more likely to be enriched further along in the pseudotime trajectories of both prototypical and proliferative lineages, as expected due to a greater replenishment post transient inflammation and were less likely to be present in the pro-disease LpM lineage. Macrophages from vehicle treated mice were most abundant in the pro-disease lineage and predominant across all pseudotime points indicating increased pro-disease differentiation (Fig.5B). Conversely, in RRx-001 treated mice, macrophages were present at earlier pseudotime points in the prototypical and proliferative lineages and were also less present in the pro-disease LpM lineage, suggesting a reduction in pro-disease phenotypical differentiation. These data led us to conclude that RRx-001 appears to prevent the acquisition of pro-disease phenotype. As our analysis revealed pseudo temporal differences at early pseudotime points, we performed an aggregate analysis of the LpM and SpM populations from RRx-001 and vehicle mice which also indicated differences in SpM gene expression (Fig.5C). To further probe these differences, we performed gene set enrichment analysis with ClusterProfiler using GO terms (RRx-001 vs Vehicle), and identified terms related to activation of macrophages, antigen processing and defence responses (Fig.5D). Furthermore, KEGG pathways suggested an enhancement of phagosome and lysosomal activity, alongside antigen processing, with the majority activated in both the LpM and SpM populations (Fig. 5E), supporting the concept that treatment of induced endometriosis with RRx-001 may impact peritoneal myeloid populations early in the differentiation trajectory. In line with RRx-001 enhancing phagocytic activity in macrophage populations we demonstrated reduced average expression level of Sirpα in transcriptomic and gene activity data (inferred from ATAC) compared to vehicle treated mice, corresponding with our flow cytometric analysis at day 14 (Figs.5F, Fig.1H) and indicating that RRx-001 acts to disrupt the Sirpα-CD47 immune evasion signal that is enhanced in endometriosis.

**Figure 5.**
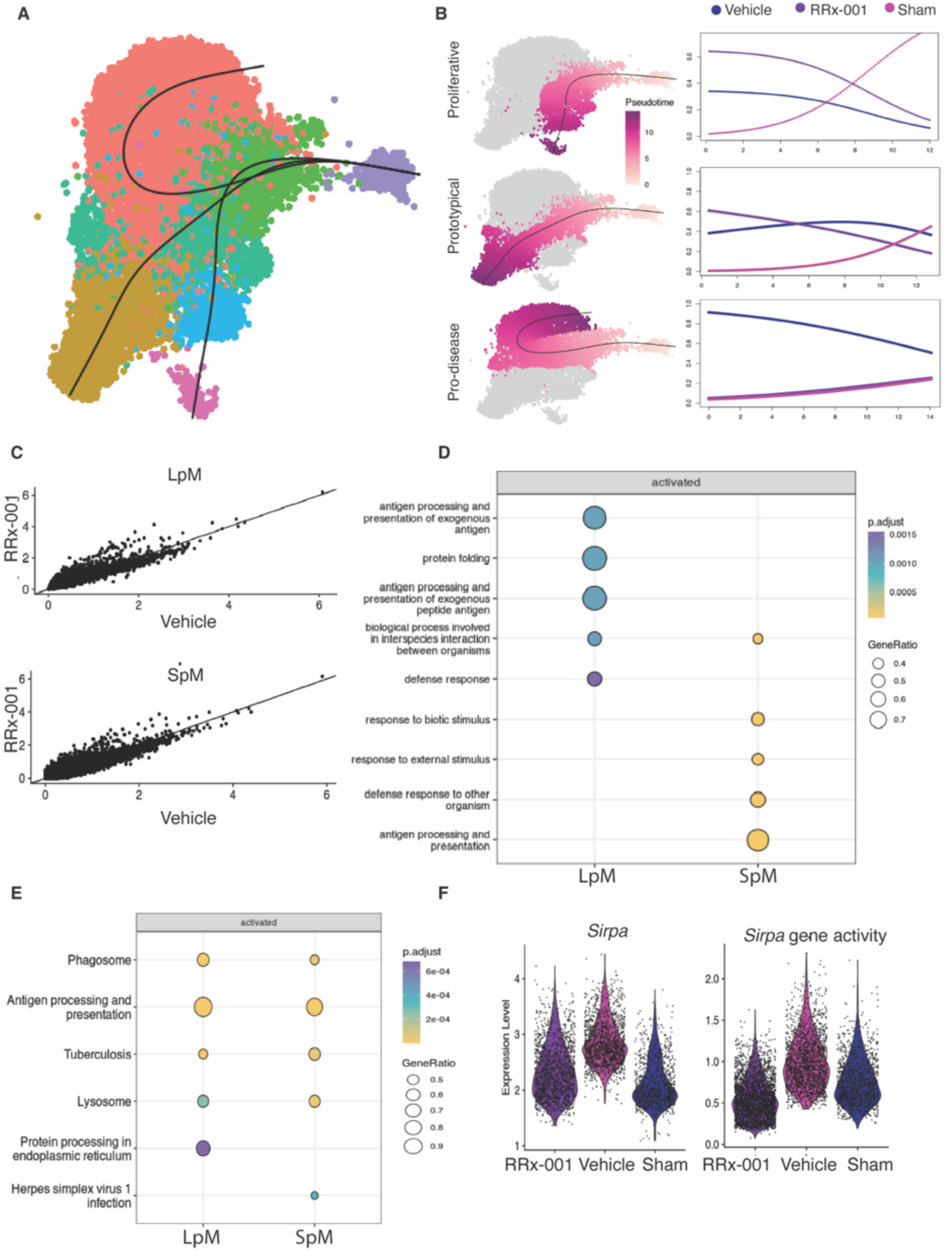
RRx-001 impacts differentiation trajectories and phenotypes of LpM and SpM populations in mice with induced endometriosis. (A) Slingshot fitted trajectories and principal curves. (B) Pseudotime ordering of inferred lineages (left) with multinomial logistic regression analysis of pseudo-temporal sample imbalance (right). (C) Scatter plot of aggregate (pseudobulk) expression of LpM and SpM populations from RRx-001 and vehicle treated mice (Pearson correlation reported). (D) ClusterProfiler gene set enrichment analysis using GO terms showing the top 5 terms in the LpM and SpM of RRx-001 treated mice with induced endometriosis. (E) ClusterProfiler gene set enrichment analysis using KEGG pathways showing the top 5 pathways in the LpM and SpM of RRx-001 treated mice with induced endometriosis. (F) Violin plots of *Sirpα* expression in RNA-seq data and the gene activity matrix derived from ATAC-seq data.

### The impacts of RRx-001 are specific to the endometriosis niche and peripheral blood

10x single cell transcriptomic profiling was used to investigate the transcriptomic impacts of RRx-001 on all Cd45+ leukocytes isolated from the peripheral blood, endometrium, liver, lungs and spleen, to determine peripheral immunological impacts and assess potential off-target effects in distal organs (Fig.6A). Fluorescent activated cell sorting (FACs) was utilised to isolate Cd45+ leukocytes from RRx-001 and vehicle treated mice in the ‘menses’ model of induced endometriosis, at day 14 post-lesion induction, followed by processing of sorted cells using the 10x genomics chromium single cell platform to generate transcriptomic libraries (n=4, n=4, respectively, Figs. 6A, S4A). An integrative UMAP visualisation of matched samples from each tissue from RRx-001 and vehicle treated was performed to enable visualisation of all 10909 cells (post quality control), revealing detection of all expected leukocyte populations as per known marker gene expression, and some organ specific localisation of cells (Fig.6B, 6C). Analysis of complexity score (log10GenesPerUMI) revealed consistency of sequencing between both organs and condition(Fig. 6D).

**Figure 6.**
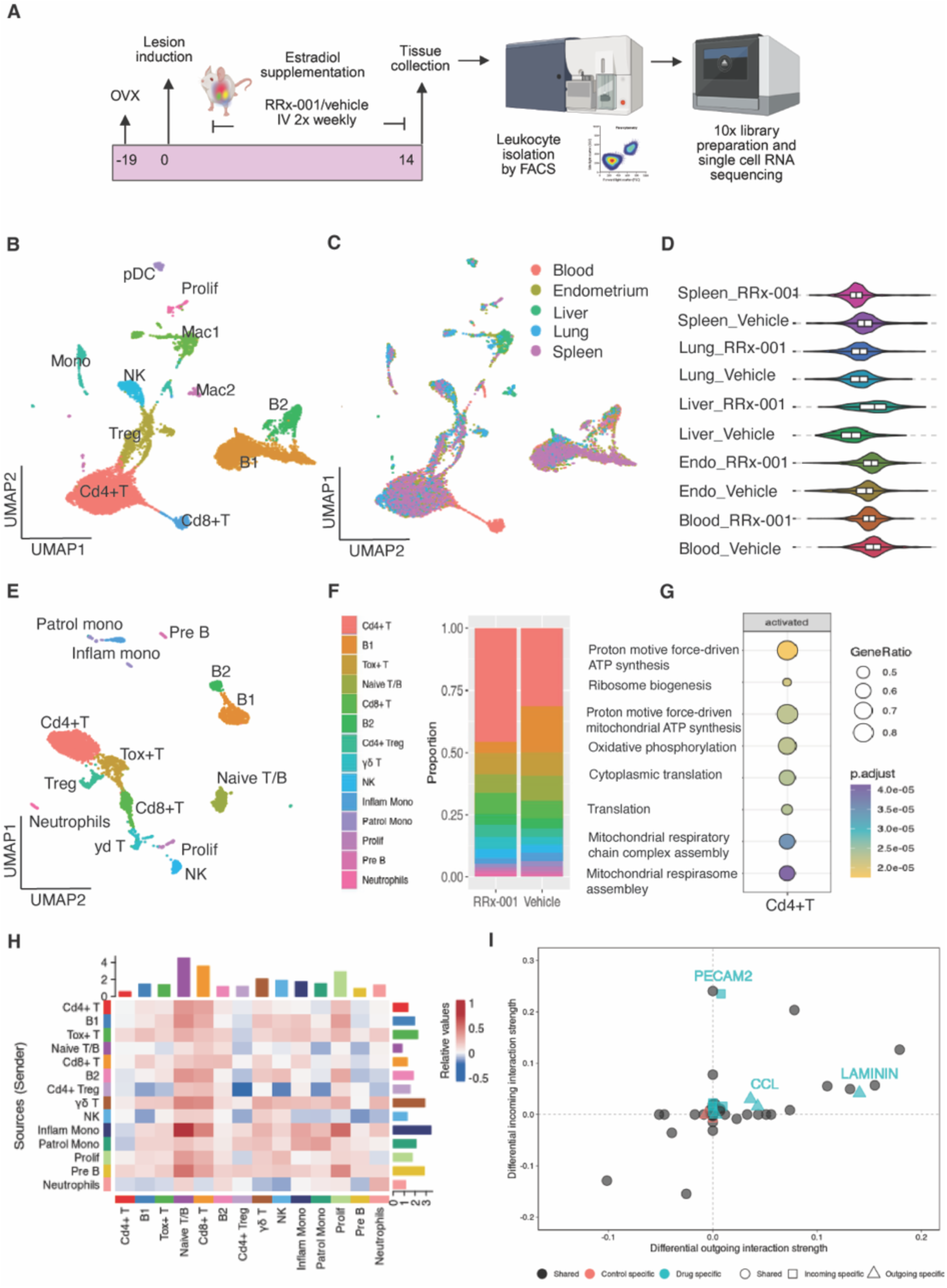
RRx-001 modifies leukocyte immune composition and cell-cell communication in the peripheral blood; distal organs are unaffected. (A) Schematic of experimental design. (B) UMAP visualisation of single cell RNA sequencing data of Cd45+ leukocytes isolated from the peripheral blood, endometrium, liver, lungs and spleen from vehicle and RRx-001 treated endometriosis mice. (C) UMAP split by tissue of origin, (D) violin plot of log10GenesPerUMI from each sample. (E) UMAP visualisation of single cell RNA sequencing data of Cd45+ leukocytes from the peripheral blood only. (F) Peripheral blood samples split by cluster composition. (G) ClusterProfiler gene set enrichment analysis using GO terms showing the top terms activated in the Cd4+ T cells from the peripheral blood of RRx-001 treated mice. (H) CellChat differential interaction strength in the peripheral blood cell-cell communication network between RRx-001 and vehicle treated mice, with red and blue representing increased and decreased signalling, respectively, in RRx-001 treated mice. (I) CellChat signalling changes of inflammatory monocytes in the peripheral blood in RRx-001 and vehicle treated mice.

We initially focussed our analysis on the peripheral blood, analysing samples from RRx-001 and vehicle treated mice. Leukocyte populations identified according to canonical markers in the blood included T cells (Cd4+ T, Tox+ T, Tregs, Cd8+ T, ψ8 T), natural killer cells (NK), B cells (B1, B2, and Pre-B), naïve T/B cells, monocytes (inflammatory monocytes and patrolling monocytes) and a proliferative cluster of immune cells (Fig.6E). An analysis of sample composition by cluster revealed all immune populations were shared between experimental groups, with an increased abundance of Cd4+ T cells expressing Ms4a6b, IL7R (naïve T cell marker) and IL6ra detected in RRx-001 treated mice (Fig.6F). We performed gene set enrichment analysis with ClusterProfiler using GO terms (RRx-001 vs Vehicle) for these expanded Cd4+ T cells, and identified terms related to metabolism, cytoplasmic translation and oxidative phosphorylation, suggesting a metabolic activation of this population (Fig.6G). Monocytes were of low abundance within the peripheral blood in both vehicle and RRx-001 treated mice; a trend of reduced Mrc1 (Cd206) was seen in patrolling monocytes following RRx-001 treatment, as previously reported in small cell carcinoma patients (Fig.6F, S4B)(*51*). We utilised CellChat to infer communication networks within the peripheral blood in RRx-001 and vehicle treated mice, and to probe condition specific signalling. CellChat also identified differential interaction strength in the cell-cell communication network among peripheral leukocytes in RRx-001 treated mice, revealing the highest differential outgoing signalling in inflammatory monocytes (Fig.6H). Probing the specific signalling changes of inflammatory monocytes by comparing the outgoing and incoming interactions, identified changes uniquely associated with RRx-001 treated mice in PECAM2, CCL and LAMININ signalling, associated with trans-endothelial migration and concurrent with our observations of increased monocyte-derived macrophages in the peritoneal fluid of these mice (Fig 6I).

To investigate any potential macrophage modulation in distal organs, we performed similar matched analyses of leukocytes isolated from the endometrium, liver, lungs and spleen. In the endometrium, UMAP visualisation was utilised for identification of leukocyte populations according to canonical markers, revealing predominant macrophages (Mac/Mono, Mac/AP and Prolif Mac) and NK cells (NK1/Cd4+ T cells, NK2 and NK3) (Fig.7A). All cell types were present in RRx-001 and vehicle treated mice, with no impacts to immune cell composition detected (Fig.7B). We utilised differential gene analysis (MAST) to analyse up and down-regulated genes in the Mac/Mono in RRx-001 compared to vehicle control mice (Fig.7C) and minimal changes in gene expression were detected (<10 highlighted DEGs), suggesting that RRx-001 does not modify endometrial macrophages. We repeated these analyses in the liver, lungs and spleen. Immune cell composition was not altered across all distal organs analysed and minimal modification of macrophage gene expression was observed (<5 DEGs) (Fig.S4C-E).

**Figure 7.**
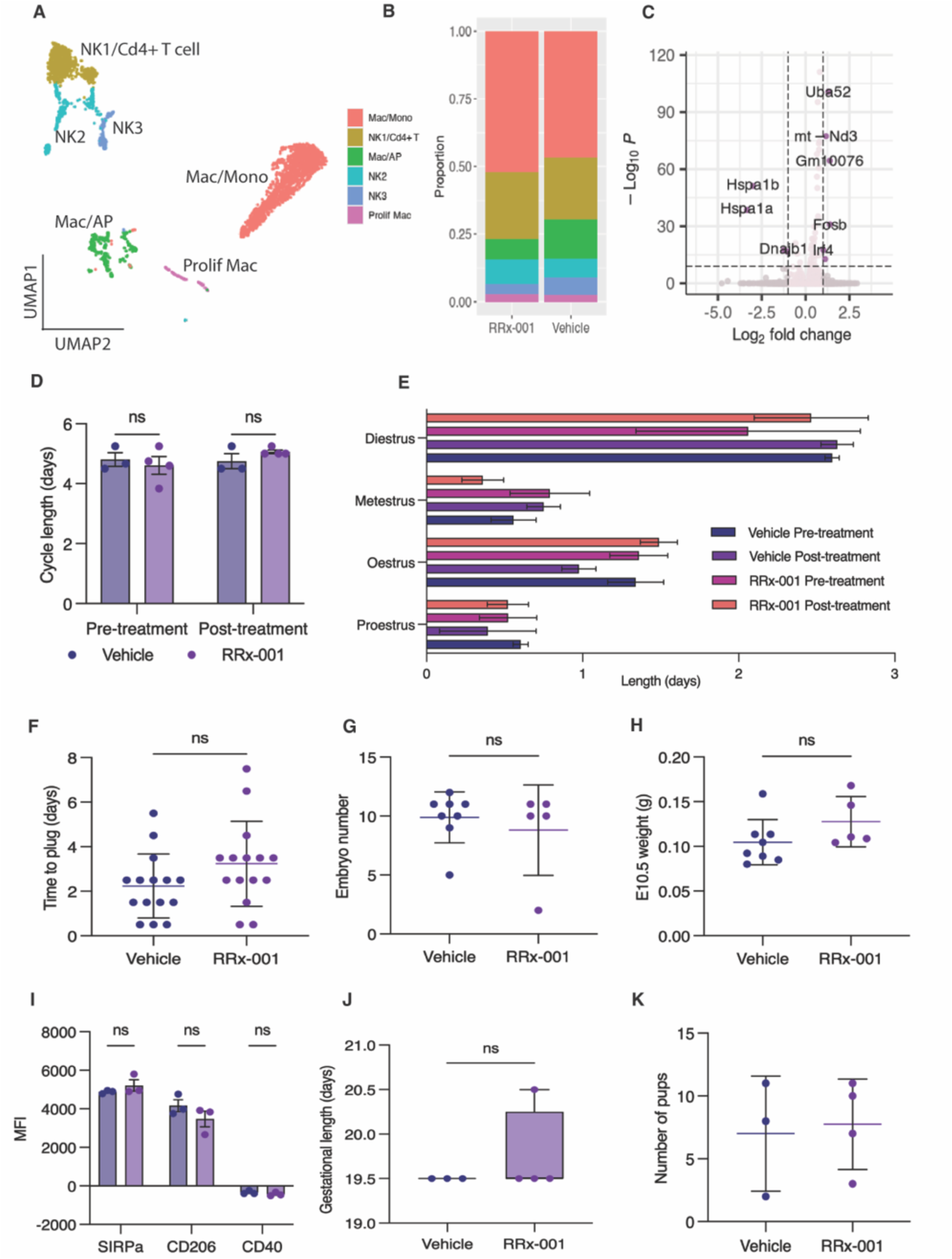
Pre-pregnancy treatment with RRx-001 does not modify endometrial macrophages or impact reproductive function. (A) UMAP visualisation of single cell RNA sequencing data of Cd45+ leukocytes from the endometrium. (B) Endometrium datasets split by cluster composition. (C) Volcano plot of up and down-regulated DEGs in the Mac/Mono endometrial population between RRx-001 and vehicle treated endometriosis mice. (D) Length of oestrous cycle and (E) days spent within each stage; n=3 vehicle, n=4 RRx-001; statistical analyses performed using Student’s t test and two-way ANOVA with Bonferroni post hoc. (F) Time to detection of seminal plug; n=15 RRx-001, n=15 vehicle; statistical analysis performed using a Student’s t test. (G) Number of embryos and (H) average weight of embryos at E10.5; n=5 RRx-001, n=8 vehicle; statistical analyses performed using a Student’s t test. (I) Median fluorescence intensity of SIRPα, CD206 (pro-repair) and CD40 (pro-inflammatory) in Hofbauer cells pooled from 3 placentas per mouse at embryonic day 15.5; n=3 RRx-001, n=4 vehicle, statistical analysis performed using Student’s t test. (J) Gestational length of pregnancy and (K) number of pups born to each mouse; n =4 RRx-001, n=3 vehicle. Statistical analyses performed using Student’s t tests. No statistical significance reached p < 0.05.

### RRx-001 does not impact reproductive function in mice

Our transcriptomic analysis revealed minimal detectable impacts to the immune cell composition in distal tissues or modification of macrophages in the endometrium and other organs. Moreover, the therapeutic activity of RRx-001 has been safely evaluated in over 300 patients with no increase in infections or anaemia detected, indicating limited side-effects. Impact on reproductive function has not yet been evaluated(*37*). Currently, gold-standard treatment of endometriosis focuses on surgical intervention or therapeutic ovarian suppression (oral contraceptives/GnRH agonists). A non-hormonal medical alternative that does not interrupt reproductive function is urgently needed. To this end, we sought to determine if pre-pregnancy treatment with RRx-001 led to any aberrant changes to fertility or developmental impacts in mice. Pre-pregnancy intravenous dosing was chosen for these initial experiments as an RRx-001-bound haemoglobin adduct persists for the lifetime of the red blood cell, reported to be 40.7 ± 1.9 (S.D) days in mice, potentially leading to long-term impacts(*52*). Tissue-resident endometrial macrophages dynamically regulate tissue homeostasis and remodelling during the oestrous cycle in mice(*53*), thus we initially investigated the impact of RRx-001 on oestrous cycle length, and within individual stages, to identify any perturbations. Oestrous cycles were synchronised using the Whitten effect(*54*), followed by daily staging by wet vaginal cytology preparations for > 4 complete cycles prior to treatment with either RRx-001 or vehicle for 1-week. Daily staging continued for > 4 further cycles. No significant changes in overall cycle length or individual cycle stage were detected in RRx-001 treated mice pre-or post-treatment (Fig.7D, E).

Next, mice were dosed with RRx-001 or vehicle for 1-week, with harem breeding set up 24-hours post final drug dose for investigations into reproductive function. Within these experiments, pregnancy success rates following copulatory plug detection were as expected within breeding colonies: 13/16 RRx-001 treated mice became pregnant (1 did not plug, 2 not pregnant post-plug) and 14/15 vehicle treated mice became pregnant (1 not pregnant post-plug). Sexual receptivity was assessed by daily seminal plug checks, detected between 0.5 - 4.5 days post-mating. No significant differences in time to seminal plug were observed (Fig.7F). At embryonic day (E)10.5, no significant differences in average embryo number or embryo weight were determined between experimental groups (Fig.7G,H). At E15.5, Hofbauer cells, a heterogenous population of placental macrophages which broadly exhibit a pro-repair phenotype, were isolated from the chorionic villus of the foetal placenta and assessed using quantitative spectral flow cytometry (Fig.S5). We sought to assess Hofbauer phenotype; dysregulation of these cells is associated with placental pathologies including inadequate development, inflammation and infection(*55*). No significant differences were identified in the MFIs of SIRPα, CD206 and CD40 between RRx-001 and vehicle groups (Fig.7I). As expected for a predominantly pro-repair population, CD206 was expressed across both experimental groups, with no expression of pro-inflammatory CD40.

Finally, the remaining pregnant mice and pups were monitored throughout pregnancy and during the postnatal period. We found no impact of RRx-001 on gestational length (Fig.7J). Furthermore, mice treated with RRx-001 did not have any alterations to number of pups (ranging from 2-11 in the vehicle group, and 3-11 in the RRx-001 treated group; Fig.7K), or sex ratio of litter (Fig.S5B). Post parturition, mothers remained healthy and omental milk spots were observed in all pups on day 1 indicative of successful lactation and nursing. Pups were weighed at postnatal day 7, with weekly weighing up to 28 days of age, with no significant differences detected at any timepoint (Fig.S5C). Collectively, these data indicate that RRx-001 does not impact reproductive function.

## Discussion

In the current study we have demonstrated that RRx-001 has a robust therapeutic impact in mouse models of induced endometriosis; treatment decreased lesion number and size, and attenuated pain associated behaviours. Our macrophage depletion study confirmed that the predominant mechanism of action was via macrophages and that they are essential to mediate the therapeutic effect. In line with this, using multiomic single nuclei sequencing we showed that RRx-001 impacted peritoneal macrophage populations whereby the abundance of pro-disease macrophages was reduced, and the proportion of pro-resolving macrophages was enhanced. Trajectory analysis inferred that RRx-001 tips the balance in favour of pro-resolving macrophage by preventing differentiation to a pro-disease phenotype. Macrophage phenotype was not modified in other tissues, indicting specificity for the endometriotic niche. Finally, preliminary investigations into reproductive function suggest that RRx-001 does not impede fertility or aberrantly affect embryonic development *in utero.* Taken together, we present promising preclinical data highlighting RRx-001 as a novel macrophage directed therapy for the treatment of endometriosis.

In the ‘menses’ model of induced endometriosis we found that when treatment with RRx-001 was initiated prior to establishment of lesions, there was no difference in bioluminescent signal between vehicle control and drug treated mice on day 3, indicating that RRx-001 does not impact the initial stages of ectopic endometrial tissue adhesion within the peritoneal cavity. This aligns with the proposed erythrophagoimmunotherapeutic activity of RRx-001; in which the covalent encapsulation of RRx-001 in red blood cells vectors RRx-001 to hypoxic (tumour) vasculature – leading to vasculature occlusion, haemolysis and erythrophagocytosis by tumour associated macrophages(*40*). This phenomenon may also be occurring in lesions, which also exhibit hypoxic areas, following the initial stages of vascularisation that occur in the first week(*56*). Interestingly, in the menses model, RRx-001 treatment led to a reduction in bioluminescent signal between days 3 to 7 and days 3 to 14, but not day 3 to 10, this may be related to cases seen in the treatment of lung cancer with RRx-001, when tumour pseudo-progression is detected early in treatment due to tumoral lymphocyte infiltration, before regression occurs later, leading us to hypothesise that this may also be occurring within the endometriotic lesions(*56*). Accordingly, when we investigated the impact of RRx-001 on established lesions in the ‘menses’ model, we identified a reduction in bioluminescent signal whereas, co-treatment of RRx-001 with liposomal clodronate (to deplete peritoneal macrophages) led to an increase in bioluminescent signal. This highlights macrophages are crucial for the therapeutic efficacy of RRx-001. Future studies should establish the dynamics of pseudo-progression, if present, and determine the regime required for complete elimination of lesions.

We also investigated the impact of RRx-001 in a minimally invasive model of induced endometriosis. We previously determined that endometriotic lesions established using this model exhibit greater longevity(*38*). In this model, full thickness uterine fragments are transferred containing myometrial tissue, resulting lesions are usually larger than in the menses model and have a cystic appearance. We used two models in this study to represent different ‘phenotypes’ of endometriosis, with the minimally invasive model representing a more persistent form of endometriosis, and following the suggestions in the WERF EPHect guidelines for experimental models that encourage the use of multiple models in preclinical testing (*41*). Here, we also demonstrated a significant reduction in lesion growth when treatment was initiated 1-week post-lesion induction, rather that prior to lesion induction, again indicative that RRx-001 resolves established lesions. In both models, treatment with RRx-001 led to an attenuation of abdominal mechanical hyperalgesia, indicative of reduced pain in this region and perhaps due to the accompanying resolution of lesions with treatment. We also included non-evoked behavioural tests as they better recapitulate impacts to well-being that are associated with endometriosis and are used as surrogate measures of visceral pain. The inclusion of non-evoked behavioural tests were described recently as an absolute requirement in the replication and translation of the complex pain experience of endometriosis patients(*43*). In our studies, nest building(*57*) was the most sensitive assay for identifying behavioural phenotypes; impaired nest building was seen in mice at day 7 post-lesion induction. Following treatment with RRx-001 nest building was significantly improved beyond that observed in naïve, pre-endometriosis mice.

Depletion of macrophages by clodronate prevented RRx-001 from impacting endometriotic lesions, demonstrating a critical role of macrophages in the anti-endometriosis activity of RRx-001. This is consistent with data from oncology studies, in where RRx-001 has been shown to modulate TAMs towards a pro-inflammatory state, with enhanced immune infiltration and vascular remodelling. The presence of infiltrating macrophages was identified as a *sine qua non* condition for the anti-tumour activity of RRx-001, with depletion of TAMs attenuating the impacts of RRx-001 in an A549 xenograft mouse model(*58*). This mechanism of macrophage modulation was reflected in the multiomic analysis, highlighting that RRx-001 leads to a reduction in phenotypically pro-disease LpM and an accumulation of a phenotypically polarised LpM, termed pro-resolving LpM (prototypical LpM remained constant in each group (sham, RRx-001, vehicle). Trajectory analysis suggested that the pro-resolving LpM phenotype is a transitory cell state, thus we predict that RRx-001 prevents the acquisition of a pro-disease phenotype by newly recruited monocyte-derived LpM, and enhances the continuous recruitment of monocytes to the peritoneal cavity. Our transcriptomic analysis supported this hypothesis, identifying differential cell-cell communication in peripheral blood inflammatory monocytes related to trans-endothelial migration following treatment with RRx-001, supporting the hypothesis of increased recruitment and extravasation of circulatory monocytes into the peritoneal cavity. This builds on previous observations of peripheral monocyte modulation in small cell carcinoma patients receiving RRx-001, such as reduced expression of pro-tumorigenic CD206 (murine equivalent Mrc1), with the same trend also detected in patrolling monocytes in mice that received RRx-001 within this current study (*51*). In small cell carcinoma patients, downregulation of CD206 was positively correlated with increased treatment response and survival, with our novel observation of enhanced extravasation and recruitment of monocytes adding to the mechanistic understanding of beneficial RRx-001 activity (*51*).

The transcriptional phenotype of pro-resolving LpM exhibited some similarities to that of monocyte-derived LpM observed in our previous studies(*26*) which were attributed to protective functions in the mouse model of endometriosis. DEGs include lipid associated genes and those encoding C1q; suggesting roles in the reduction of available lipids, hormone synthesis and activation of membrane lytic complexes. In patients with endometriosis, lipidomic studies have highlighted dysregulated lipid metabolism and elevated levels of certain lipid species, possibly driven by oxidation stress and inflammation in the body, suggesting a potentially beneficial effect of enhancing lipid metabolism and warranting further investigation in endometriosis research (*59, 60*). C1q, the initiator of the classical complement pathway, places a multifaceted role in immune surveillance, inflammation and tissue remodelling, processes all relevant to the pathophysiology of endometriosis. We observed enhanced C1q following RRx-001 treatment, which may promote the clearance of ectopic endometrial cells by facilitating the opsonization and removal of apoptotic cells and their cellular debris, while concurrently suppressing inflammasome activation, thus controlling peritoneal inflammation. Similar protective effects have been reported in early atherosclerotic lesions, a disease which shares many commonalities in the involvement of macrophages and inflammatory pathways; complement activation via C1q has been identified as protective in early lesions by dampening inflammation, benefits which diminish at later disease stages (*61, 62*). Concurrently, the pro-angiogenic and immune-evasive properties of C1q may contribute to pathological angiogenesis and persistence of established lesions at later timepoints, indicating its role may be time-dependent in the progression of the disease and suggesting the need for temporally informed immune modulation strategies in endometriosis (*63*). Treatment with RRx-001 also appeared to increase genes associated with response to pathogens and antigens, again suggesting improved visibility of ectopic tissue to peritoneal macrophages and potentially enhanced clearance of apoptotic debris. Together these modifications to peritoneal macrophages may reduce endometriosis drivers and promote destruction of ectopic tissue.

RRx-001 has previously been shown to have significant impacts on the epigenome of cancer cells; effects vary depending on the dose and timepoints evaluated. Sites that were hypo- and hyper-methylated as well as globally increased acetylation were reported(*64*). We predict that in the peritoneal cavity RRx-001 prevents acquisition of pro-disease phenotype via epigenetic modification. Of particular interest, we found that Klf (including Klf14) binding motifs were enriched in peritoneal macrophages derived from mice exposed to RRx-001. Previous studies have identified a role for Klf14 in reducing glucose metabolism and immune function in macrophages; this seems to exert a protective function in inflammatory conditions by limiting inflammatory cytokine secretion(*49*). Further studies should validate these findings mechanistically. These modifications also highlight the paradoxical nature of RRx-001 mechanisms reported in the literature, whereby antigenicity is enhanced whilst simultaneously promoting an antioxidative and anti-inflammatory environment (*65*).

Much of the research conducted on the mechanism of action of RRx-001 has occurred in the oncology field, where studies describe targeting of RRx-001 to hypoxic and aberrant vasculature prevalent within tumours via binding to haemoglobin within red blood cells. Drug bound RBCs travel to hypoxic vasculature where they accumulate because of their RRx-001 induced rigidity(*66*). During this process, nitric oxide (NO) is released which is thought to confer cytotoxic activity within tumours(*67*). Macrophages also engulf RRx-001 bound RBCs, inducing polarisation from a pro-repair to pro-inflammatory state and down-regulating the CD47-SIRPα pathway, which reduces tumour cell immune evasion(*68, 69*). Immunogenicity of cancer cells is also induced via increased expression of interferon (IFN)-responsive genes(*70*). Given the similarities that exist between cancer and endometriosis (including pro-disease macrophage polarisation and pathological immunotolerance) and that endometriosis lesions also exhibit a hypoxic environment, we predict that significant RRx-001 impacts are also occurring within lesions. In support of this, the transcriptomic analyses performed in this study infer such a targeted mechanism, as we detected profound immunological changes only in the peritoneal fluid with a fine tuning of the systemic immunity identified in peripheral blood. Future work will interrogate the intra-lesional environment in response to RRx-001. In the current study, we failed to evaluate this as the lesions recovered were so reduced in size that we were unable to recover enough cells for scRNA-Seq studies on multiple occasions. The mediation of indirect effects within associated environments has not yet been characterized, however from our results it is plausible to suggest that the modulation of macrophage populations within the peritoneal cavity occurs as a secondary outcome resulting from endometriotic lesion resolution by lesion-resident macrophages and a subsequent reduction in the signals that dictate the acquisition of a pro-disease LpM phenotype, or by reactivating a previously immunotolerant microenvironment within the peritoneal cavity. The release of reactive oxygen and nitrogen species may also play a role. Thus, further work is required to elucidate the impacts of RRx-001 on lesion-resident macrophage populations.

To investigate potential impacts on fertility, we evaluated the endometrium immune environment and reproductive parameters in mice exposed to RRx-001. Our findings suggest that RRx-001 does not impact the immune cell composition within the endometrium or modify endometrial macrophage gene expression. Additionally, pre-pregnancy treatment with RRx-001 does not impair the normal reproductive function of female mice or embryo development indicating that it has potential suitability as a non-hormonal and fertility sparing treatment for endometriosis. Further work will identify whether RRx-001 can rescue endometriosis-associated infertility.

To summarize, we have revealed a novel indication for the anti-cancer drug RRx-001 in the treatment of endometriosis. It acts to reduce endometriosis lesions, associated pain-like behaviours and modulates macrophage populations in the peritoneal cavity without negatively impacting fertility or modifying macrophages in distal tissues. RRx-001 is currently in phase I - III clinical trials for different solid cancers, is well tolerated with available safety data. In this first phase of preclinical testing, we highlight a robust effect in experimental endometriosis and plan to proceed to a feasibility trial in women with the disorder. RRx-001 and its macrophage modifying actions has the potential to improve the lives of millions of women suffering from endometriosis.

## Acknowledgements.

The authors thank Dr Laura Baxter at the Bioinformatics RTP, University of Warwick, for pre-processing of multiomic data; the Biological Services Unit, University of Warwick, for care of mice; and the FlowSRL, University of Warwick, for assistance with flow cytometry experiments. We would also like to thank Deborah McIntyre for the independent quantitative scoring of nests.

## Author contributions

E.G conceptualised and supervised the study and acquired funding. I.M conducted all experiments, analysed data, and performed multiomic and transcriptomic analyses. V.V performed multiomic analyses. C.L.P and M.R performed experiments. P.D and K.P performed animal experiments. I.M and E.G wrote the manuscript.

## Data and materials availability

All data associated with this study is present within the manuscript and the included supplementary materials. Analysis script for multiomic and transcriptomic analyses are publicly available on github.com under project tbc. Multiomic sequencing data are deposited in Gene Expression Omnibus under accession number tbc.

## Materials and Methods

### Study design

Experiments were designed to determine the impact of RRx-001 treatment in a mouse model of experimental endometriosis and to identify any potential impact on fertility outcomes. For the purpose of determining the impact of RRx-001 on induced endometriosis, we implemented two complementary mouse models (‘menses’(*71*) and ‘minimally-invasive’(*38*)) in which to robustly examine endometriotic lesion growth and pain-like behaviours. A macrophage depletion study in the ‘menses’ model allowed us to confirm the effects seen were mediated by peritoneal macrophages. Additionally, we utilised multiomic sequencing to determine transcriptomic and epigenomic effects of RRx-001 on peritoneal macrophages isolated from the ‘menses’ mouse model, and single cell transcriptomic sequencing of Cd45+ leukocytes isolated from the peritoneal fluid, peripheral blood and distal organs (liver, lung, spleen and endometrium) to assess cell-cell communication and potential off-target impacts. To assess fertility outcomes, we examined the effect of pre-pregnancy dosing of RRx-001 on the oestrous cycle and reproductive functions. Animals were randomised to either RRx-001 treatment or control groups using the GraphPad QuickCalcs online tool (https://www.graphpad.com/quickcalcs/randomize1/). Behavioural experiments were performed under blinded conditions. Details of mouse models, sample size and statistical methods are described in the appropriate sections below.

### Animals

Wild-type FVB/N mice were purchased from either Envigo (Cambridge, UK) or Charles River (UK) and maintained under standard conditions at the University of Warwick, UK. FVB-Tg(CAG-luc,-GFP)L2G85Chco/J (stock number 008450|L2G85) were originally purchased from The Jackson Laboratory (Bar Harbor, ME, USA), and a breeding colony maintained under standard conditions at the University of Warwick. Mice were housed in standard conditions with a 12:12 light-dark cycle, an ambient temperature of 21°C, humidity at 50%, and *ad libitum* access to food (standard rodent chow) and water. The diet of weanlings was supplemented by apple sauce. Following arrival to the facility, mice acclimatised for one week prior to commencement of experimental procedures. All procedures performed were permitted under license by the United Kingdom Home Office Animal Experimentation (Scientific Procedures) Act; PPL PP0568394, and in accordance with the guidelines of the Committee for Research and Ethical Issues of the International Association for the Study of Pain.

### Mouse models of experimental endometriosis

We implemented two variations of the bioluminescent syngeneic mouse models of induced endometriosis; a ‘menses’ model and a minimally invasive model, described in detail previously(*38, 71*). In brief, for the ‘menses’ model, recipient FVB/N mice (n=40) were ovariectomised and received an intraperitoneal injection of decidualised and progesterone withdrawn donor endometrium (*72*) recovered from FVB/Cag-Luc mice (∼40mg, minced, combined, resuspended in saline and passed through an 18 g needle; n = 40 FVB/Cag-Luc donors) and exogenous oestradiol supplementation (oestradiol valerate 500 ng, twice weekly). Surgical controls, referred to as ‘sham’ mice, were ovariectomised FVB/N mice that received intraperitoneal injection of saline instead of tissue. In the minimally invasive model, intact recipient FVB/N mice (n = 32) received an intraperitoneal injection of FVB/Cag-Luc donor full-thickness uterus (equivalent to 1 uterine horn per recipient, minced and passed through an 18 g needle; n = 16 FVB/Cag-Luc donors). Oestrus stage was determined in donors and recipients by vaginal smear and retrospective cytological analysis(*73*).

### Experimental groups

Mice were randomly assigned to either (I) RRx-001 treatment, (II) vehicle control, (III) RRx-001+clodronate group, (IV) sham or (V) naïve (no procedures). Experimental groups included in each experiment are as tabulated below. During these experiments, 5 mice were culled early due to adverse effects (venous irritation of tail) and were not included in reported numbers or data. Dosing schedules are noted in individual experimental schematics. In all our studies, RRx-001 (Selleck Chemicals, USA) was dosed intravenously at 10mg/kg in DMA:PEG-400 1:2, diluted 1:4 in sterile saline immediately prior to injection, 2x weekly (Monday and Thursday, 12pm), with control mice receiving vehicle (DMA:PEG-400 1:2) only.

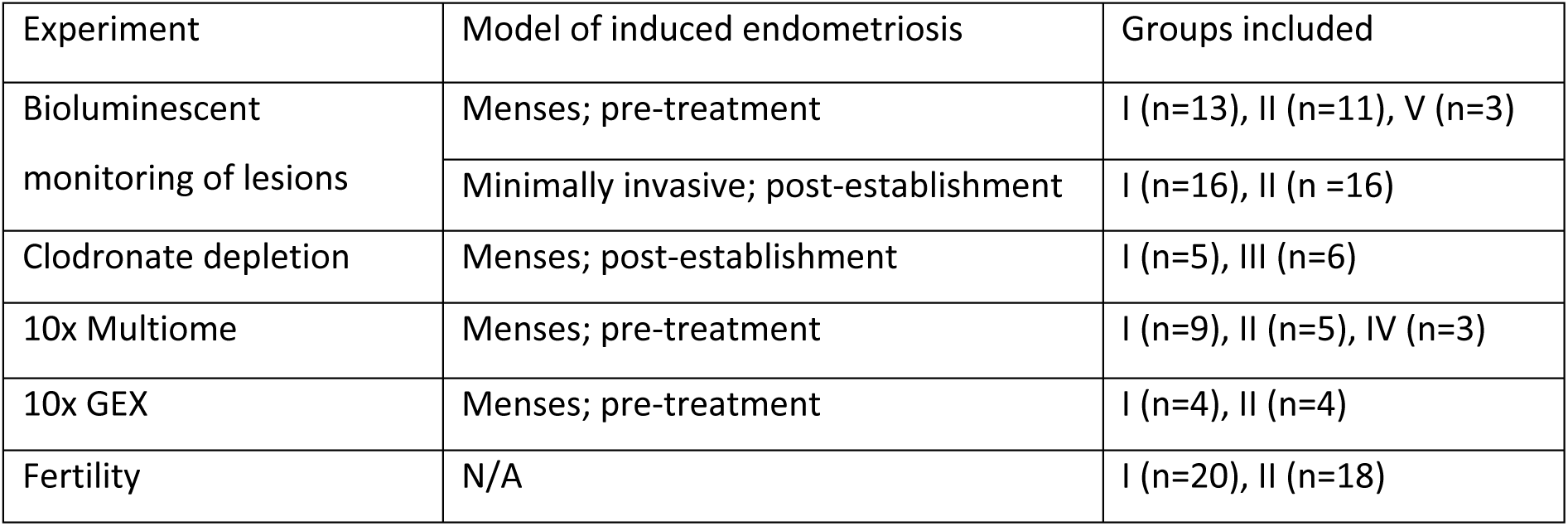

### Non-invasive *in vivo* bioluminescence quantification

As described previously(*39*), mice were injected subcutaneously with 1.5 mg D-Luciferin Potassium Salt (75mg/kg, reconstituted in sterile PBS and sterilised by 0.22 μm filtration), following a 5-minute period to allow a maximal luciferase signal plateau, mice were imaged under maintained anaesthesia using a PhotonIMAGER (Biospace Lab, Paris)(*38*). Lesion bioluminescence was measured using a standard region of interest across experiments, on the front and back of each mouse for 7 minutes per side. Using M3 Vision software (Biospace Lab), the background reading was subtracted, front and back readings averaged, and photons/s/cm2/sr calculated for bioluminescent signal quantification.

### Histological and ImageJ analyses

Post tissue fixation and embedding in paraffin, 3 μM sections of recovered lesions were stained using a standard haematoxylin/eosin protocol. Images were taken on a LeicaDMI microscope (Leica, Germany), with white balance and histological verification of endometriotic lesion hallmarks (stroma +/-glandular epithelium) conducted in FIJI (ImageJ2, Version 2.3.0). In FIJI, analysis to determine cross-sectional area was performed on lesion images captured using 4x magnification, image scale was set to a known size and the ‘select’ tool was implemented to outline the lesion while excluding peritoneal wall/adipose tissue, and the ‘measure’ function was then used to calculate cross-sectional lesion area.

### Behavioural assessments

#### von Frey

We determined mechanical allodynia as an average of measurements taken on days 11/12/13 post lesion induction (‘menses’ model) or day −1 (baseline), 8 (1 week; initiation of treatment) and 18 (10 days following start of treatment; minimally invasive model) using calibrated Semmes-Weinstein von Frey filaments (Stoelting, Wood Vale, IL, USA), as described previously(*38, 74*). Briefly, mice were allowed to adapt to the apparatus until exploratory responses ceased. Filaments were then applied 10 times in ascending order to the abdomen or hind paw. The filament force (grams) was recorded when withdrawal responses were observed in 50% of applications.

#### Hot plate

Thermal hyperalgesia was determined prior to lesion induction, and on day 7 and day 17 post-lesion induction using a hot plate (Ugo Basile, Italy). Mice were positioned onto a preheated plate set at 52°C and their responses were observed. A time-to response (seconds) was noted when mice exhibited a reaction to the thermal stimulus. If no response was observed within 25 seconds, the mice were removed from the plate.

#### Thermal plate preference

Thermal hyperalgesia was determined prior to lesion induction, and on day 7 and day 17 post-lesion induction using TPP thermal place preference plates (Ugo Basile, Italy). The thermal plates were set at distinct temperatures, one at 12.5°C and the other at 30°C. Mice were allowed a 15-minute-period for exploration of both plates, during which the duration spent at 12.5°C was recorded.

#### Burrowing

The ethological rodent behaviour of burrowing was determined prior to lesion induction and on day 8 and day 18 post-lesion induction. Mice were individually supplied with a burrowing tube (diameter 6.8 cm, length 20 cm), made of clear acrylic, open and elevated by 3 cm at one end to prevent the accidental displacement of burrowing substrate (tubes manufactured at Life Sciences, University of Warwick). The mouse was placed in a plastic cage with the burrowing tube filled with 110g of wooden shredding; following a one-hour burrowing assay, the material remaining was weighed and recorded, and the percentage of material displaced within an hour calculated.

#### Nesting

Nesting (also ethological) was determined prior to lesion induction and on day 8 and day 18 post-lesion induction as described(*57*). Briefly, 1 hour prior to dark phase, mice were individually housed with wooden shredding and a Nestlet (VetTech, UK). The following morning, the nest was imaged, and the quality of the nest created from the shredded nestlet recorded with a rating scale of 1-5. (e.g. 1 no nest; 5 fully shredded material arranged into a neat nest). Mice had *ad libitum* access to food and water during the test and were re-grouped immediately following the nesting assay (Table S1).

### Flow cytometry

Peritoneal lavage was collected into 5 ml of ice-cold DMEM, red blood cells were lysed (Red blood cell lysis buffer, Miltenyi Biotech) and cells filtered through a 40 μM filter and then blocked with monocyte block (TrueStain Monocyte Blocker™ BioLegend), prior to resuspension and staining of 1 million cells per sample with the antibody panel shown. Samples were analysed using a FACS Aria Fusion with FACS Chorus software (BD Biosciences) and FlowJo v.9 software. Hofbauer cells were isolated from the foetal placenta of mouse embryos at E15.5. Dissection of the foetal placenta from embryonic yolk sacs and subsequent placental digestion into a single cell suspension were performed as described previously, with three foetal placentas pooled from each mouse(*75, 76*). Samples were filtered, RBCs lysed, blocked, and resuspended as above, prior to staining of 1 million cells per sample with the antibody panel. Samples were analysed on a Sony spectral ID7000 spectral cytometer (Sony Biotechnology, USA).

| Marker | Fluorophore | Clone | Supplier | Product Number |
| --- | --- | --- | --- | --- |
| CD45 | Pacific Blue | 30-F11 | Biolegend | 103125 |
| Sirpα | BV510 | P84 | Biolegend | 144005 |
| Ly6C | BV711 | HK1.4 | Biolegend | 128037 |
| CD40 | FITC | 3/23 | Biolegend | 124607 |
| c-kit | PerCP Cy5.5 | ACK2 | Biolegend | 135133 |
| CD163 | PE | S15049F | Biolegend | 156703 |
| CD34 | PE-Cy5 | MEC14.7 | Biolegend | 119311 |
| CD206 | PE-Cy7 | C068C2 | Biolegend | 141719 |
| Lineage exclusion | APC |  | Biolegend |  |
| CD19 |  | 1D3/CD19 |  | 152410 |
| CD3 |  | 17A2 |  | 100235 |
| CD335 |  | 29A1.4 |  | 137607 |
| Ly6G |  | 1A8 |  | 127613 |
| SiglecF |  | S17007L |  | 155507 |
| MHCII | APC-Cy7 | M5/114.15.2 | Biolegend | 107627 |
| F4/80 | APC Fire750 | BM8 | Biolegend | 123117 |
| Live/Dead | Zombie UV | - | Biolegend | 423107 |

### Preparation of peritoneal immune cells for 10X multiomic sequencing

Peritoneal lavage was pooled within experimental groups (endometriosis/RRx-001 n=9, endometriosis/vehicle n=5, and sham n=3) prior to processing and staining as described above, with fluorescence activated cell sorting based on

surface marker expression (Lin-/Cd45+) performed to enrich samples for macrophage populations. 100,000 cells per sample were sorted into 1ml PBS + 2% FBS using a FACs Aria. Gentle nuclei isolation was performed, and nuclei were then processed and barcoded according to the established pipelines to generate individual paired RNA and ATAC libraries using a 10x Nuclei Isolation Kit, a 10x Multiome Kit and 10x Chromium Controller^TM^ (10x Genomics). Libraries were sequenced individually by the Genomics Lab at the University of Warwick on high-output 150 cycle cartridges using a NextSeq 500/550 (Illumina®, San Diego, USA).

### Preparation of leukocytes for 10X single cell sequencing

All samples were pooled within experimental groups (endometriosis/RRx-001 n=4 and endometriosis/vehicle n=4). Peritoneal fluid was processed and stained as described above. Peripheral blood was collected via cardiac puncture, prior to processing and staining as above. Mechanical disruption was used to prepare the spleen, prior to processing and staining as above. Endometrial, liver, and lung tissues were digested for 20 minutes, 45 minutes, and 60 minutes, respectively, in 3 ml of a solution containing Collagenase IV (0.2 mg/mL) and DNase I (0.05 mg/mL) in RPMI1640 at 37°C, prior to processing and staining as above. Fluorescence activated cell sorting based on surface marker expression (Cd45+) was then performed to isolate leukocytes. 100,000 cells per sample were sorted into 1ml PBS + 2% FBS using a FACs Aria. Sorted cells were then processed using a 10x Chromium Next GEM Single Cell 3’ Kit v3.1, and the 10x Chromium Controller^TM^ (10x Genomics) to generate transcriptomic libraries. Libraries were sequenced by Novogene (Novogene Co., Ltd).

### Bioinformatic analysis of peritoneal multiomic data

We followed recent guideline best practices for single cell analysis across modalities as described by Heumos et al for our paired analysis of snRNA-seq and snATAC-seq libraries(*77*). BCL files for each flow cell directory from the ATAC and GEX sequencing runs were demultiplexed using Cell Ranger ARC analysis pipelines to generate FASTQ files. Seurat (5.0.3) and Signac (1.13.0) in R studio (4.3.2) were used to create Seurat objects (v3) based on GEX data, with ATAC-seq data added as a second assay. Using ATAC-seq data, a common peak set was created and peaks were called in individual samples using MACS2 v2, with peaks on non-standard chromosomes and in genomic blacklist regions removed(*78*). Independent quality filtering of each dataset was performed on both modalities to remove low-quality cells based on number of detected molecules for each modality, the number of genes detected per UMI, nucleosome signal and TSS enrichment. As described in datasets generated by Yoshimura et al and 10x Genomics, the gentle nuclei isolation protocol allows mitochondria to attach to the nuclei prior to GEM generation resulting in a high mitochondrial percentage(*79*). Post-matrix generation we filtered reads aligned to mitochondrial or ribosomal regions, and matrices were then subset on common genes. Finally, the scDblFinder and SoupX were used to identify and remove doublets and assess ambient contamination, respectively(*80, 81*). Pre-processing of RNA datasets was performed using the SCTransform v1 workflow (Seurat), followed by conservative anchor-based RPCA-integration using the default k.anchor parameter (5) to align populations (Seurat). We then computed the LSI of individual peak datasets using standard functionalities (Signac), followed by Harmony integration of datasets(*82*). The Seurat v4 weighted nearest neighbour (WNN) method of ‘FindMultiModalNeighbors’ was utilised to compute a joint neighbour graph to represent both the GEX and ATAC modalities, followed by the use of standard workflows for WNN UMAP visualisation and clustering (Seurat). Clusters were identified using marker genes identified in Wilcoxon rank-sum tests in a one-vs-all fashion for manual annotation according to our previously published annotations of mouse peritoneal macrophages(*26*). Clusters corresponding to non-macrophage populations or expressing < 1% of total cells were removed in order to isolate 11,877 peritoneal macrophages (Fig.S3B, C). To account for within-sample correlation we implemented MAST, a generalised mixed effect model, for differential gene expression testing(*83*). Differentially accessible peaks were identified utilising logistic regression within the Signac ‘FindMarkers’ function. DNA sequence motif information (JASPAR2020) was added using the ‘AddMotifs’ function, with enriched motifs identified using ‘FindMotifs’.

For data visualisation purposes the Seurat and Signac functionalities ‘DimPlot’, ‘FeaturePlot’, ‘DotPlot’, ‘CoveragePlot’, ‘CellScatter’, ‘VlnPlot’, ‘MotifPlot’, and the EnhancedVolcano package were implemented(*84*) (*85, 86*). The clusterProfiler package was used for statistical analysis, output simplification and visualisation of GO and KEGG terms(*87*). For trajectory analysis, DynGuidelines was utilised to select the most applicable package and the indicated Slingshot package were implemented(*88, 89*).

### Bioinformatic analysis of leukocyte transcriptomic data

BCL files for each flow cell directory from the GEX sequencing runs were demultiplexed using Cell Ranger analysis pipelines to generate FASTQ files. BCL files for each flow cell directory from the GEX sequencing runs were demultiplexed using Cell Ranger analysis pipelines to generate FASTQ files. Seurat (5.3.0) was utilised in R studio to create Seurat objects (v5) based on filtered feature count matrices. Independent quality filtering of each sample was performed to remove low-quality cells based on percentage of mitochondrial reads, complexity (log10GenesPerUMI > 0.8), number of genes and UMI counts. Merged objects of paired data from RRx-001 and vehicle treated mice were created in each tissue type (endometrium, blood, lungs, liver and spleen) for further analysis. Standard Seurat workflow of ‘NormalizeData’, ‘FindVariableFeatures’, ‘ScaleData’, ‘RunPCA’, ‘FindNeighbours’ and ‘FindClusters’ (SLM algorithm) was performed, with RunUMAP utilised for UMAP visualisation of clusters. Clusters with no expression of CD45+ were removed from Seurat objects. Clusters were identified and manually annotated according to marker genes identified using Wilcoxon rank-sum tests in a one-vs-all fashion. Data visualisation was performed using the Seurat ‘DimPlot’ and scPubr ‘do_ViolinPlot’ functions (*90*). For differential gene expression testing between RRx-001 and vehicle samples, we implemented MAST, a generalised mixed effect model, with data visualisation using EnhancedVolcano. Paired datasets were combined from all organs and integrated using Harmony, followed by visualisation using UMAP. The clusterProlifer and CellChat packages were utilised for statistical analysis, output simplification and visualisation of GO terms, and to infer and visualise differential cell-cell communication networks, respectively (*91*).

### Statistics

Statistical analyses were performed using GraphPad Prism 10.1.1 for macOS (San Diego, CA). All graphical data plotted as mean with standard deviation. For data meeting the necessary assumptions for parametric analyses (Shapiro-Wilk), Student’s t-tests or two-way analysis of variance (ANOVA) with post hoc testing conducted with the Bonferroni test. If non-normality of data was detected, non-parametric analyses were performed using a Mann-Whitney *U* test. Statistical significance for all tests was established at *P* < 0.05. Significance levels were denoted as follows: * *P* < 0.05, ** *P* < 0.01, *** *P* < 0.001, **** *P* < 0.0001.

## Supplementary Figures

**Figure S1:**
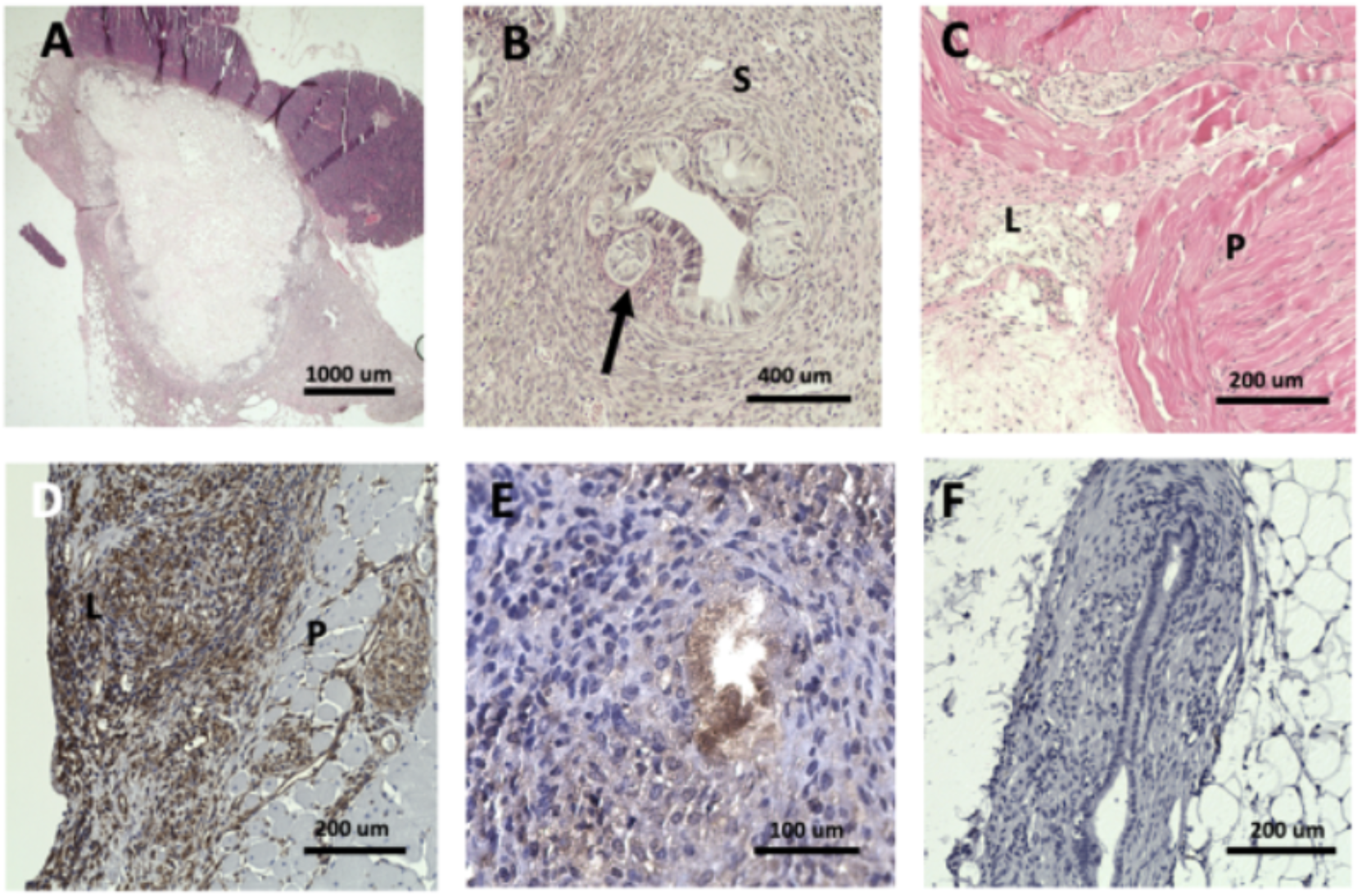
Histological verification of lesions. (A-C) Representative hematoxylin and eosin stains of recovered lesions. (A) Cystic lesion, (B) glandular (arrow) and stromal structures (S) within lesion. (C) Lesion (L) infiltration into the peritoneal wall (P). (D) Stromal (vimentin^+^ cells) on a peritoneal wall lesion. (E) Epithelial (cytokeratin^+^ cells) showing staining of a glandular structure. (F) Negative control. Scale bars as shown.

**Figure S2:**
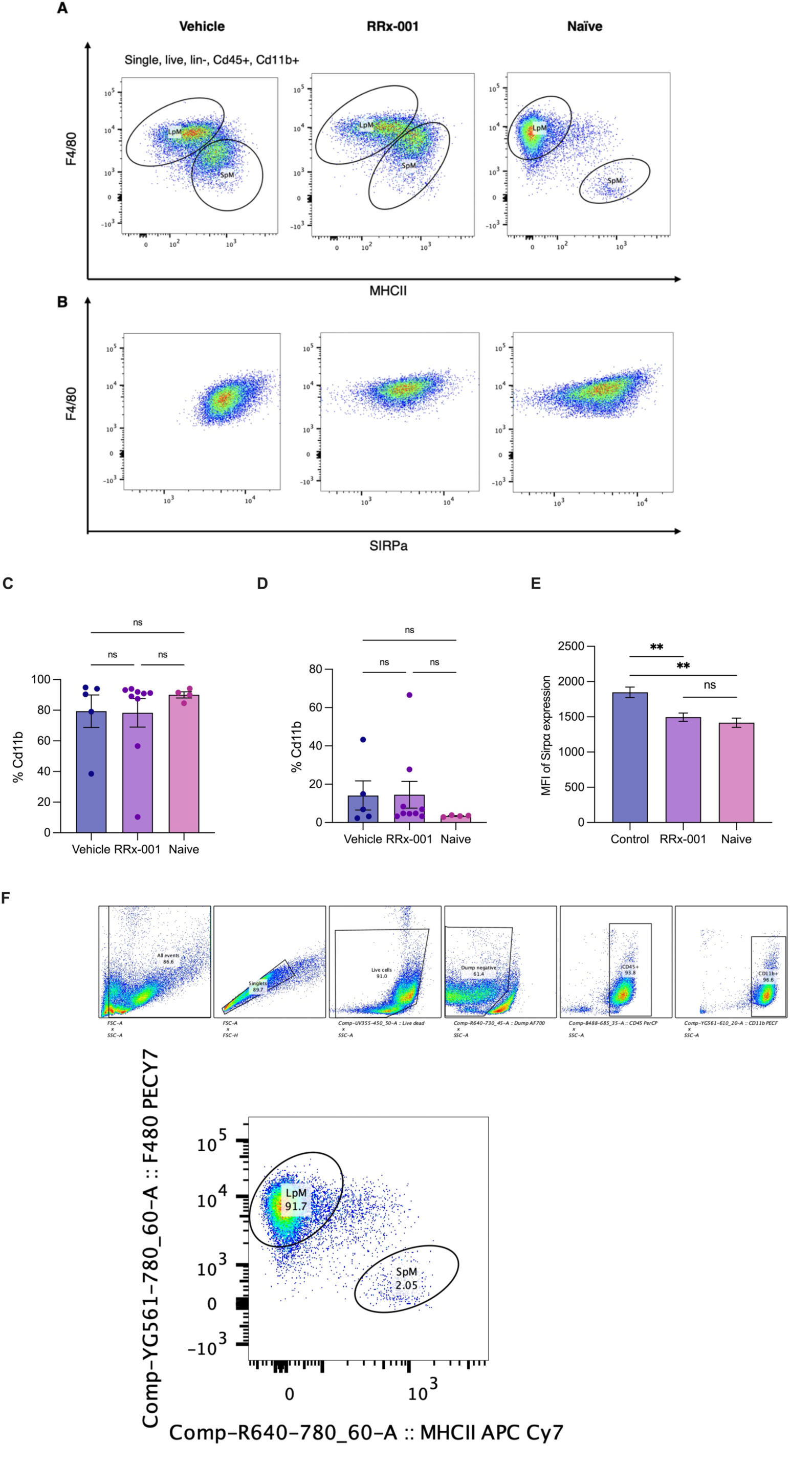
Representative flow plots of peritoneal macrophage sub-populations and expression of Sirpα. (A) Representative flow plots of LpM and SpM populations in vehicle, RRx-001 treated and naïve mice. (B) LpM population and Sirpα expression. (C) Percentage of Cd11b+ cells that are LpM and (D) SpM. (E) Median fluorescence intensity of Sirpα on LpM. (F) Flow cytometry gating strategy.

**Figure S3:**
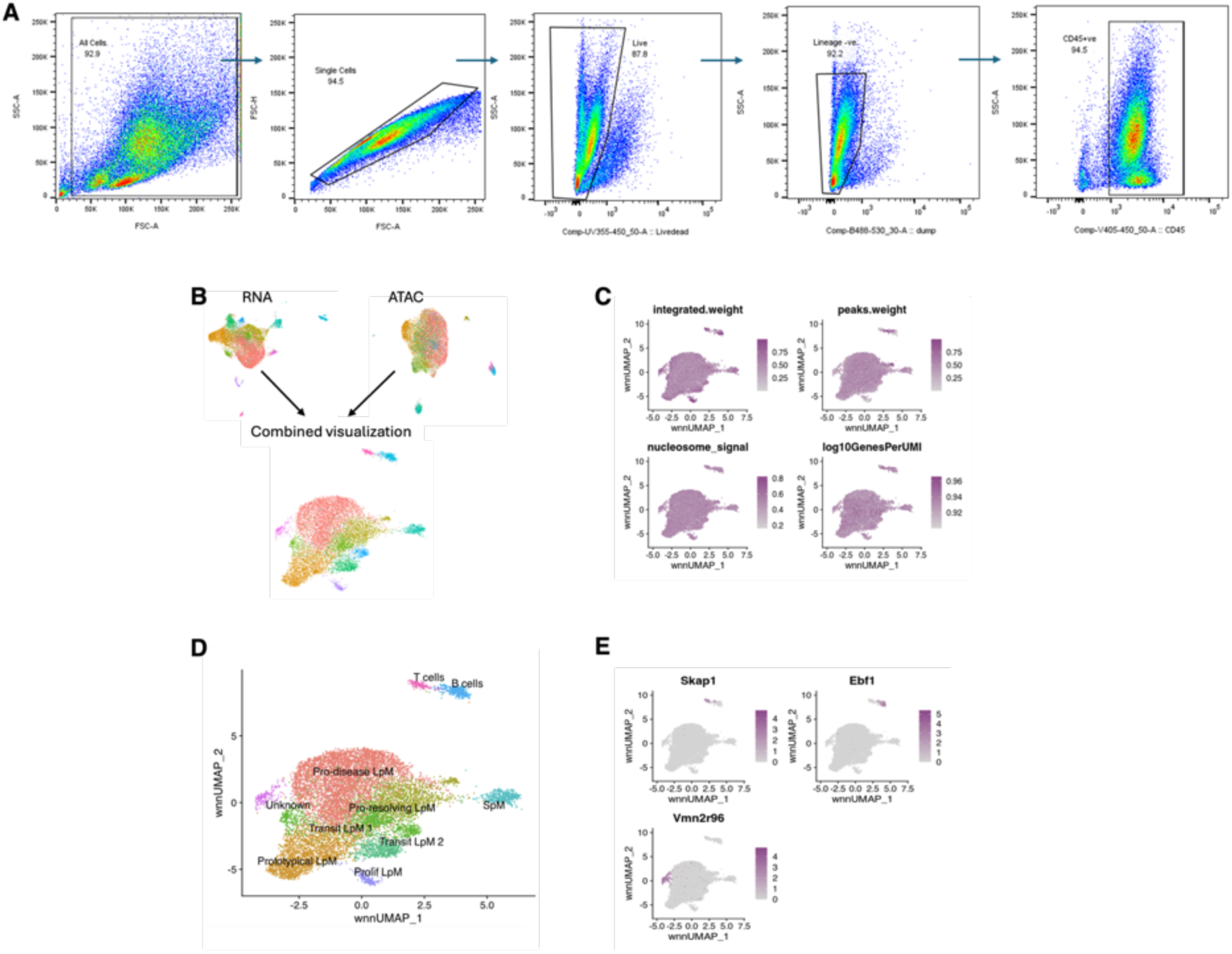
Isolation of macrophage for integrative weighted nearest neighbours (WNN) analysis. (A) Flow cytometry gating strategy. (B) UMAP visualisations of individual RNA and ATAC modalities and their integrated joint WNN visualisation. (C) RNA and ATAC modality weight across clusters, nucleosome signal (ATAC) and log10GenesPerUMI (RNA). (D) Clusters corresponding to T, B or unknown immune cells were removed by identification using marker genes shown in (E) Feature plots of Skap1, Ebf1 and Vmn2r96, respectively.

**Figure S4:**
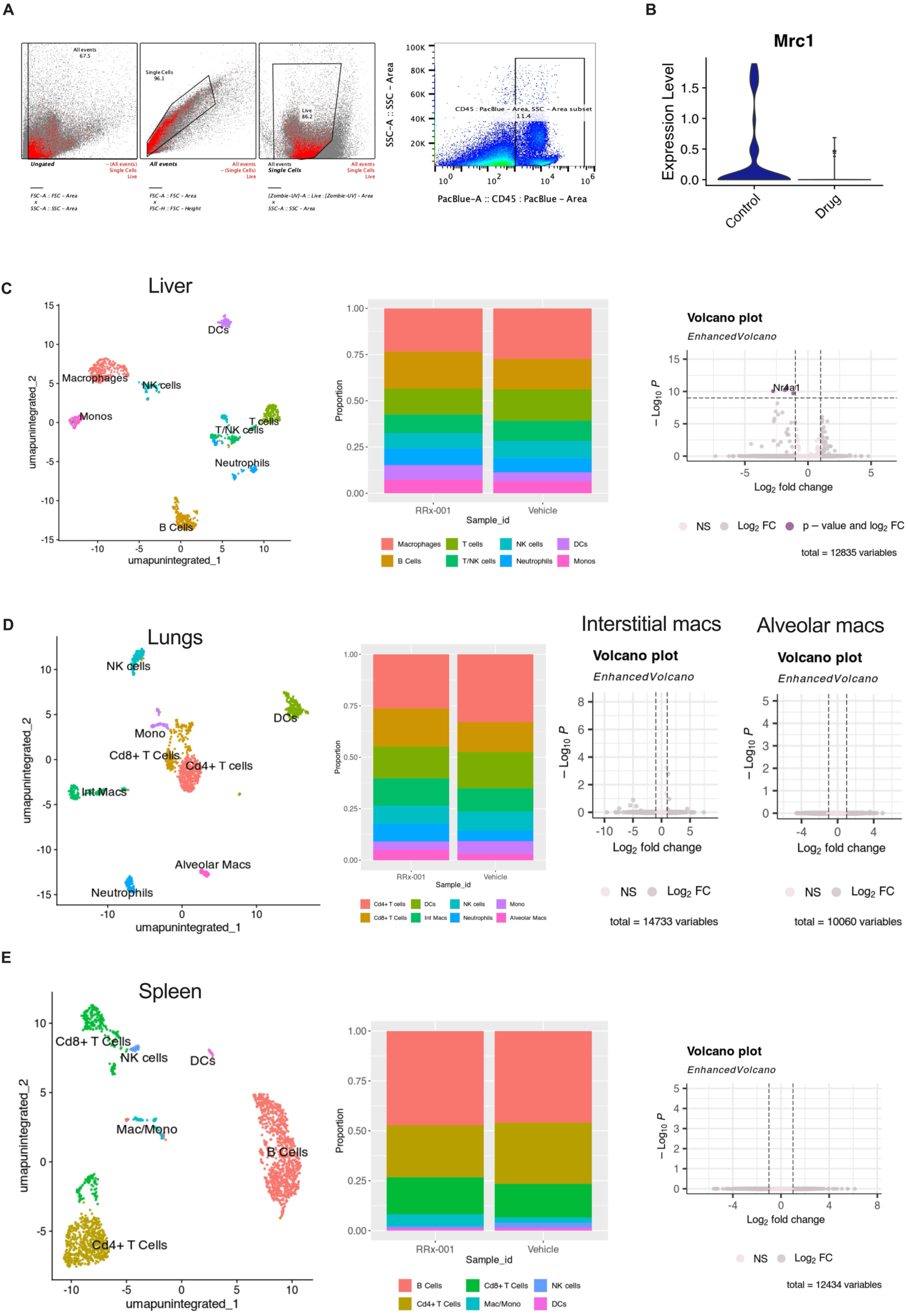
Gating strategy for isolation of CD45+ leukocytes and the analysis of distal organ immune composition and potential macrophage modification following RRx-001 treatment. (A) Flow cytometry gating strategy. (B) Violin plot of Mrc1 on patrolling monocytes. (C) UMAP of Cd45+ leukocytes isolated from the liver of RRx-001 and vehicle treated mice, cluster composition of each sample, and volcano plot of DEGs between RRx-001 and vehicle treated mice in the macrophage population. (D) UMAP of Cd45+ leukocytes isolated from the lungs of RRx-001 and vehicle treated mice, cluster composition of each sample, and volcano plots of DEGs between RRx-001 and vehicle treated mice in the interstitial and alveolar macrophage populations. (E) UMAP of Cd45+ leukocytes isolated from the spleen of RRx-001 and vehicle treated mice, cluster composition of each sample, and volcano plot of DEGs between RRx-001 and vehicle treated mice in the macrophage population.

**Figure S5:**
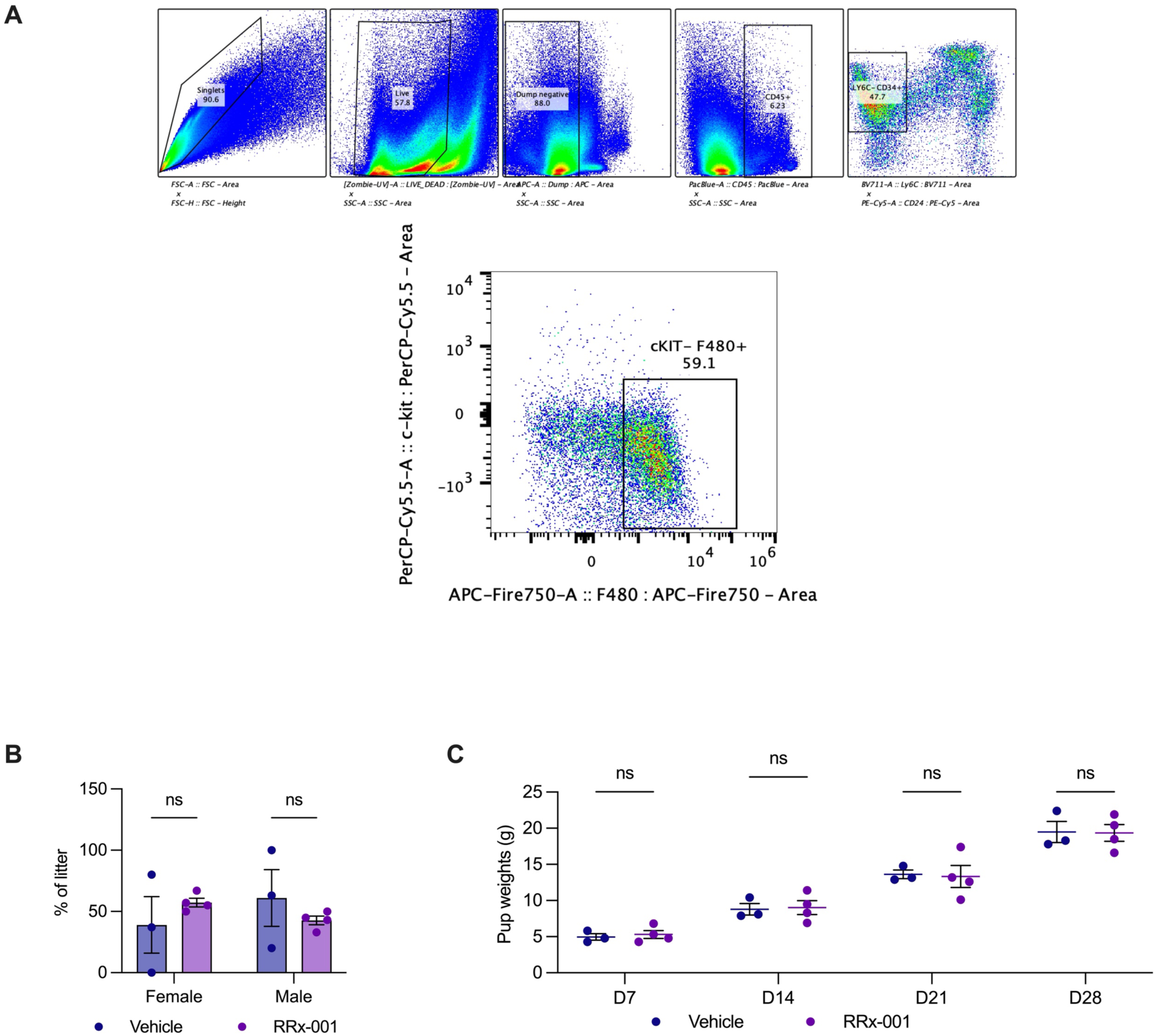
Gating strategy for isolation of Hofbauer cells and pup reproductive outcomes following pre-pregnancy treatment with RRx-001 or vehicle control. (A) Flow cytometry gating strategy. (B) sex ratio of litter and (C) average litter weights from postnatal day 7 to postnatal day 28; n=4 RRx-001, n=3 vehicle. *S*tatistical analyses performed using Student’s t tests. No statistical significance reached p < 0.05.

